# Evolution of the spider homeobox gene repertoire by tandem and whole genome duplication

**DOI:** 10.1101/2023.05.26.542232

**Authors:** Madeleine E. Aase-Remedios, Ralf Janssen, Daniel J. Leite, Lauren Sumner-Rooney, Alistair P. McGregor

## Abstract

Gene duplication generates new genetic material that can contribute to the evolution of gene regulatory networks and phenotypes. Duplicated genes can undergo subfunctionalisation to partition ancestral functions and/or neofunctionalisation to assume a new function. We previously found there had been a whole genome duplication (WGD) in an ancestor of arachnopulmonates, the lineage including spiders and scorpions but excluding other arachnids like mites, ticks, and harvestmen. This WGD was evidenced by many duplicated homeobox genes, including two Hox clusters, in spiders. However, it was unclear which homeobox paralogues originated by WGD versus smaller-scale events such as tandem duplications. Understanding this is key to determining the contribution of the WGD to arachnopulmonate genome evolution. Here we characterised the distribution of duplicated homeobox genes across eight chromosome-level spider genomes. We found that most duplicated homeobox genes in spiders are consistent with an origin by WGD. We also found two copies of conserved homeobox gene clusters, including the Hox, NK, HRO, *Irx*, and SINE clusters, in all eight species. Consistently, we observed one copy of each cluster was degenerated in terms of gene content and organisation while the other remained more intact. Focussing on the NK cluster, we found evidence for regulatory subfunctionalisation between the duplicated NK genes in the spider *Parasteatoda tepidariorum* compared to their single-copy orthologues in the harvestman *Phalangium opilio*. Our study provides new insights into the relative contributions of multiple modes of duplication to the homeobox gene repertoire during the evolution of spiders and the function of NK genes.

## INTRODUCTION

Whole genome duplications (WGDs) have occurred several times during animal evolution (Ohno 1970; Putnam et al. 2008; Flot et al. 2013; Kenny et al. 2016; Schwager et al. 2017; Nong et al. 2021). Duplicated genes are thought to be released from selective constraints, allowing paralogues to subfunctionalise, specialise, or neofunctionalise either by changes to their cis-regulatory controls or protein sequence while collectively maintaining the robustness of the ancestral functionality (Force et al. 1999; Jiménez-Delgado et al. 2009; Tinti et al. 2012; Espinosa-Cantú et al. 2015; Marlétaz et al. 2018). The two rounds of WGD (2R WGD) in the gnathostome ancestor has been credited with facilitating the evolution of vertebrate novelties, however, the extent to which WGDs in other animal lineages have contributed to diversification is not well understood (Meyer and Schartl 1999; Shimeld and Holland 2000; Cañestro et al. 2013; Simakov et al. 2020; Aase-Remedios and Ferrier 2021).

Many retained ohnologues (genes duplicated via WGD) are transcription factors that regulate processes in development, which further implicates the role of duplication in the evolution of phenotypes and the underlying developmental regulatory networks (Brunet et al. 2017). Homeobox genes are a superclass of transcription factors that expanded by tandem duplication early in animal evolution into eleven recognised classes encompassing more than 100 bilaterian gene families with diverse and wide-reaching roles in development (Table 1) (Holland et al. 2007; Larroux et al. 2008; Zhong et al. 2008; Kumar 2009; Butts et al. 2010; Bürglin 2011; Zhong and Holland 2011; Holland 2013; Holland et al. 2017). Certain homeobox gene expansions may have coincided with innovations that occurred at the evolutionary origins of animals and subsequently bilaterians (Garcia-Fernàndez 2005; Holland 2013; Holland 2015). Tandem duplication of homeobox genes early in animal evolution resulted in clusters of related genes that have been conserved across animals (Pollard and Holland 2000; Larroux et al. 2007; Larroux et al. 2008; Ferrier 2016a). Homeobox gene clusters (Table 1) are thought to be maintained by functional constraint on their genomic organisation exerted by complex and overlapping regulatory elements controlling the expression of adjacent genes (Irimia et al. 2012; Ferrier 2016b).

**Table 1.**
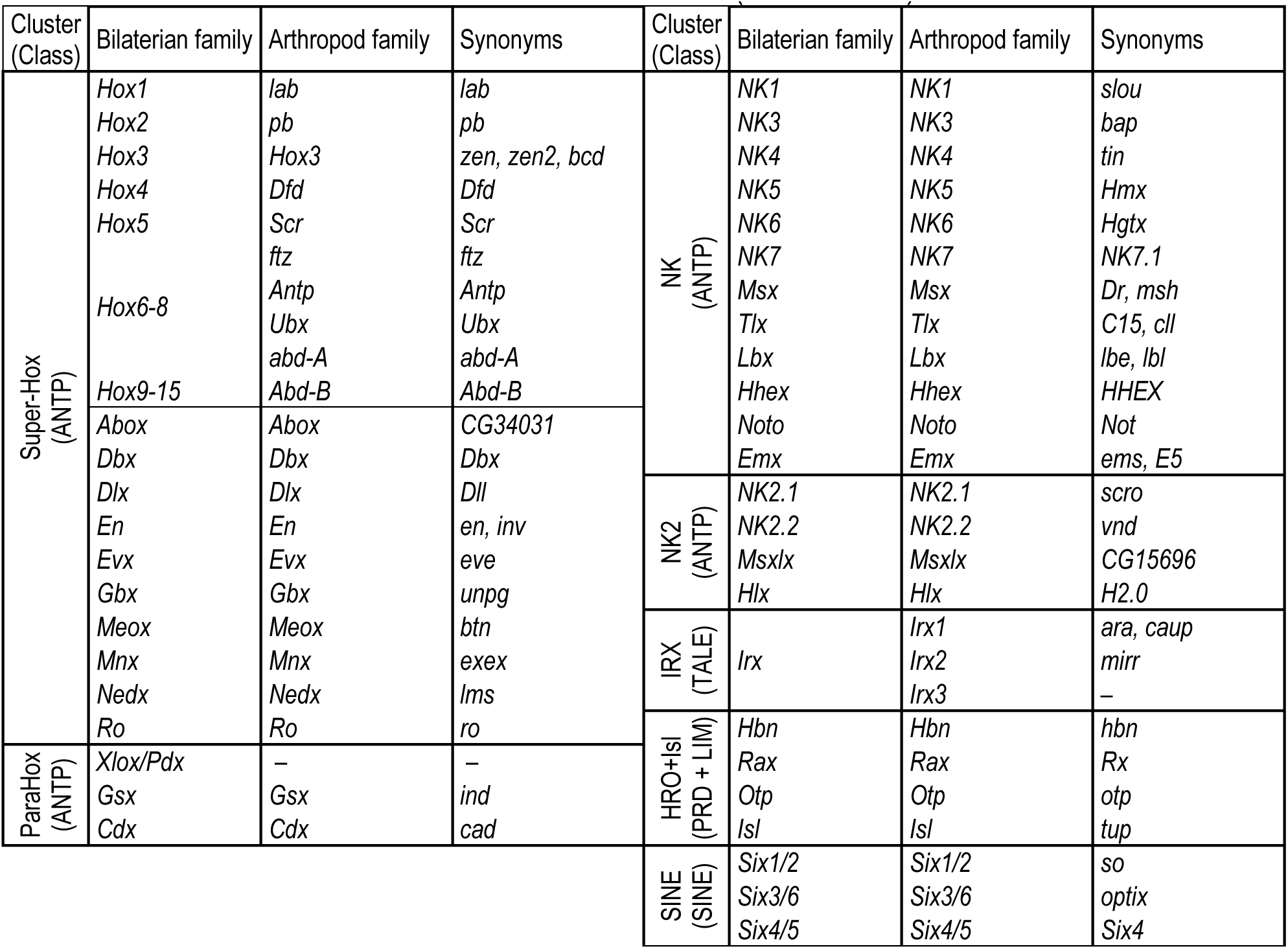
– Genes and *Drosophila melanogaster* synonyms in conserved bilaterian homeobox clusters. Bilaterian homeobox gene **families** contain genes descendant from a single inferred gene in the ancestor of all bilaterians, as previously defined by Ferrier (2016b) and the references therein. Families are grouped into **classes,** which themselves originated by duplication earlier in animal evolution. **Clusters** are composed of related genes that arose by tandem duplication and have been conserved across bilaterians. The Hox cluster contains the Hox genes, while the ancestral bilaterian Super-Hox cluster includes the related ANTP-class genes that are often found linked to the Hox cluster (Butts et al. 2010; Ferrier 2016b). The ParaHox cluster is the sister of the Hox cluster and consists of three bilaterian families; dashes for *Xlox/Pdx* indicate it is not found in arthropods (Brooke et al. 1998; Ferrier 2016a). The NK cluster also belongs to the ANTP class. Some ANTP-class NK2 cluster genes were first described as part of a ‘pharyngeal’ cluster, containing *NK2.1*, *NK2.2*, and *Msxlx*, as well as two non-homeobox genes, *Pax1/9* and *FoxA1/2* (Simakov and Kawashima 2017); linkage and sometimes clustering between *NK2.1*, *NK2.2*, and *Msxlx* is evident across bilaterians, and here is referred to as the NK2 cluster (Simakov et al. 2015; Ferrier 2016b). The PRD-class HRO cluster (Mazza et al. 2010) also includes the LIM-class gene *Isl* as they are often found linked (Ferrier (2016b). The TALE-class *Irx* cluster likely underwent independent duplications in different lineages; *ara* and *caup* are a fly-specific duplication of one *Irx* family, often clustered with single arthropod *mirr* orthologues of a second *Irx* family (Irimia et al. 2008; Kerner et al. 2009). The dash for *Irx3* indicates it is not found in *D. melanogaster*. The SINE-class cluster (Gallardo et al. (1999) consists of three families and is conserved across bilaterians (Ferrier 2016b).

The best known homeobox gene cluster, the Hox cluster, illustrates how the genomic organisation of homeobox genes is important for their function. Most arthropod genomes contain one Hox cluster consisting of ten ANTP-class genes (Table 1), which are involved in patterning the antero-posterior axis of animals in the same relative order as the genomic organisation of the genes within the cluster (Krumlauf 2018 and the references therein). The Hox cluster is highly conserved across bilaterians, both in genomic organisation and function, though differences in Hox gene expression are known to underlie changes to animal body plans (Averof and Patel 1997; Abzhanov et al. 1999; Janssen et al. 2014; Martin et al. 2016; Serano et al. 2016; Janssen and Pechmann 2023). Hox clusters have also been retained in duplicate following WGDs, as exhibited by the four vertebrate Hox clusters that arose via the 2R WGD (Holland and Garcia-Fernàndez 1996; McLysaght et al. 2002; Holland et al. 2007; Putnam et al. 2008; Cañestro et al. 2013; Holland 2013; Holland and Ocampo Daza 2018). The 2R WGD appears to have relaxed constraint on vertebrate Hox cluster organisation so that while the complement of genes from *Hox1* to *Hox14* are present in at least one copy, no cluster contains all 14 genes (Hoegg and Meyer 2005). It is still not well understood to what extent the patterns of gene loss between duplicate clusters and the subfunctionalisation observed for vertebrate homeobox gene clusters apply more widely to other homeobox gene clusters (Table 1) and to other animal lineages that have undergone WGD.

Besides gnathostomes, an independent WGD took place in the arachnopulmonate ancestor approximately 450 MYA (Figure 1), resulting in nearly 60% of homeobox genes retained in duplicate in spiders and scorpions and their relatives, including two Hox clusters (Schwager et al. 2017; Leite et al. 2018; Harper et al. 2021). Within each Hox gene family, one ohnologue exhibits different spatial and temporal expression compared to the other, though overall, spatial and temporal collinearity is largely intact for each cluster (Schwager et al. 2007; Schwager et al. 2017; Turetzek et al. 2022). Several of the other retained homeobox genes exhibit divergent expression consistent with both sub- and neofunctionalisation in *Parasteatoda tepidariorum*, showing that duplication of these genes has played a role in the evolution of spider development (Leite et al. 2018).

**Figure 1.**
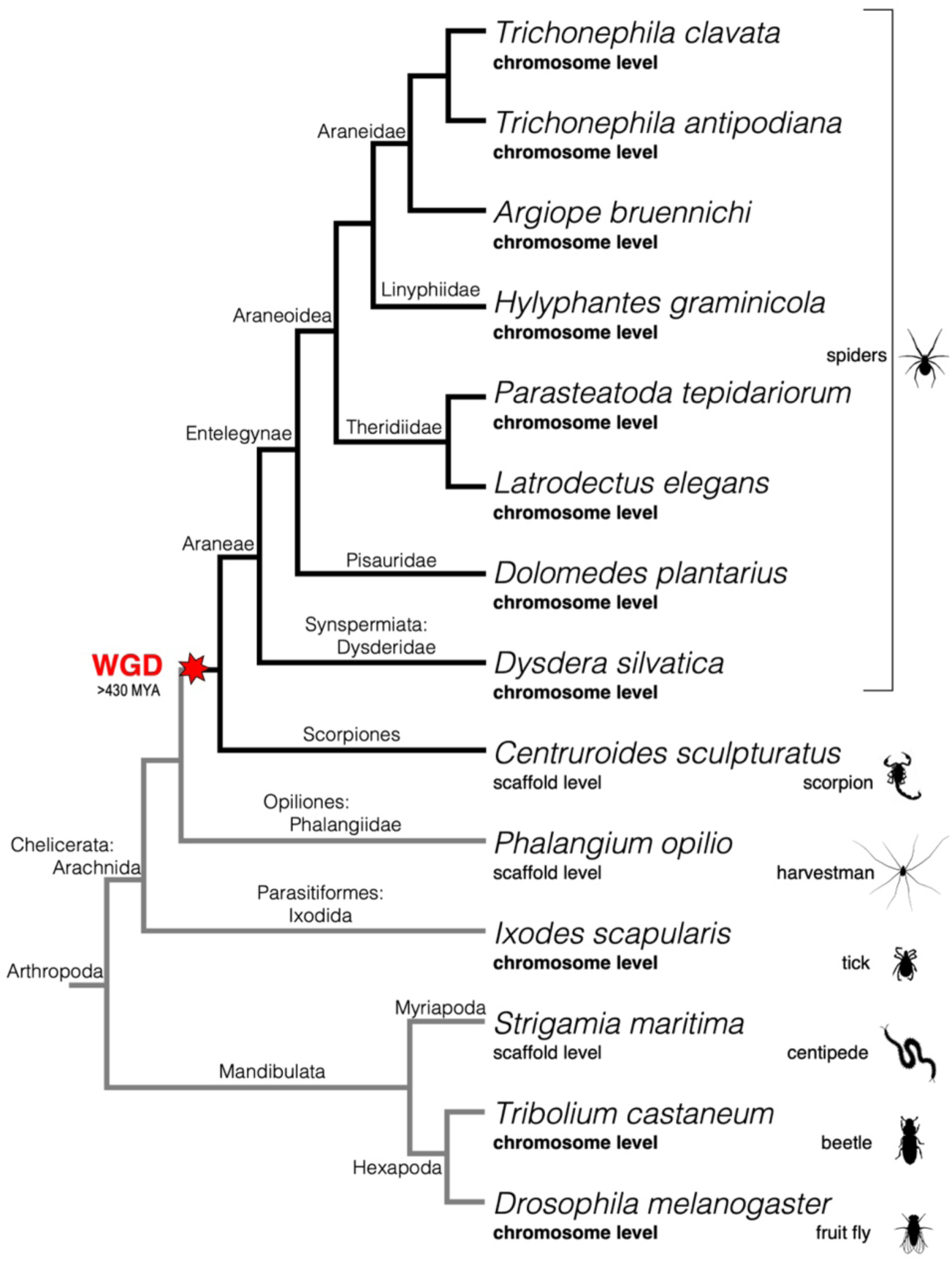
– Cladogram of arthropods used in this study. The red star indicates the timing of the whole genome duplication (WGD); pre-duplicate lineages have grey lines, duplicate lineages have black lines. Genome assembly level is indicated below the species binomial, in bold for the chromosome-level assemblies. Higher phylogenetic classifications are labelled on phylogeny nodes. The topology was drawn based on published molecular phylogenies (Ontano et al. 2021; Li et al. 2022). Silhouettes were taken from the PhyloPic database [https://www.phylopic.org/].

The lack of chromosomal-level genome assemblies for spiders has so far hindered a full understanding of the relationships among duplicated homeobox genes and the relative impact of the WGD compared to tandem duplication. It was not possible to determine whether clusters were fully intact or had undergone degradation, beyond one of the Hox clusters (Schwager et al. 2017). Leite et al. (2018) described clustering of a few genes, each belonging to several of the other homeobox gene clusters besides the Hox, including fragments of the NK, HRO, *Irx*, and SINE clusters, some of which were found in duplicate. However, it remained impossible to determine whether these duplicates were ohnologues, indicating the retention of ohnologous clusters, or if they resulted from tandem duplications occurring before or after the WGD. Furthermore, it could not be determined if these duplicates were shared with other spider species because of the lack of comparative genomic resources.

We surveyed the homeobox gene complement and synteny across eight recently released chromosome-level genome assemblies available for spiders including *P. tepidariorum* (Figure 1, see Materials and Methods). Our analysis allowed us to distinguish WGD ohnologues from paralogues that arose via tandem duplication for the first time among spiders. We found that many spider homeobox genes and clusters were retained in duplicate following the WGD, within conserved patterns of whole-genome synteny. We also characterised both ancient and recent tandem duplications and determined the timing of these relative to the WGD and the divergence of major lineages within arthropods. We found that the NK cluster was retained in duplicate and largely conserved across spiders. Complementing the previous analysis of *Emx* and *Msx* family genes (Leite et al. 2018), we described the expression of the rest of the NK cluster genes during embryogenesis in the harvestman *P. opilio*, an outgroup to the arachnopulmonates that did not undergo the WGD, and their orthologues and ohnologues from the spider *P. tepidariorum*.

For these NK cluster genes, we detected no evidence of selection on coding sequences in our comparisons between paralogues or between single copy *Phalangium opilio* genes and their *P. tepidariorum* ohnologues, but for each family with retained ohnologues in *P. tepidariorum*, we found distinct expression patterns for each ohnologue during embryogenesis, providing evidence for regulatory subfunctionalisation. Combined, these results provide new insight into both the patterns of ohnologue retention and their consequences in spider evolution.

## RESULTS

### Homeobox gene repertoires in arthropods

We surveyed the homeobox genes of eight spider species with chromosome-level genome assemblies, representing seven entelegyne and one synspermiatan species and compared these to the scorpion *Centruroides sculpturatus*, the harvestman *Phalangium opilio*, and the tick *Ixodes scapularis*, as well as three mandibulates (Figures 1, 2 and Table 2). We found that spiders and scorpions have similar numbers of homeobox genes, and on average, have 1.4 times more homeobox genes than outgroup species that did not have an ancestral WGD (Table 2). The spiders and scorpion had an average of 150 homeobox genes; the most are found in *Hylyphantes graminicola* with 172 and the fewest in *Dysdera silvatica* with 140. Mandibulates and non-arachnopulmonate arachnids had on average 107 homeobox genes, with the fewest in the insects (103) and the most in *Strigamia maritima* (113). Previous surveys found 96 and 69 homeobox genes in *I. scapularis* and *P. opilio*, respectively,145 in *P. tepidariorum*, and 156 in *C. sculpturatus* (Schwager et al. 2017; Leite et al. 2018), but we identified between 1-55% more in each of these species (Table 2). For *P. opilio*, searching both the recent genome assembly (Gainett et al. 2021) as well as the developmental transcriptome (Sharma et al. 2012) provided a more complete dataset and enabled the identification of more homeobox genes than previously described.

**Table 2.** – Homeobox genes across arthropods are largely stable, except for an increase in the arachnopulmonates. Homeobox families: the number of described bilaterian homeobox gene families with at least one gene present in a genome (Holland et al. 2007; Zhong et al. 2008; Zhong and Holland 2011; Ferrier 2016b). Tandem duplications are divided into ancient (predating the WGD) and recent (after the WGD). Species to the right of the vertical line had an ancestral WGD. Species abbreviations: *Dmel Drosophila melanogaster*, *Tcas Tribolium castaneum*, *Smar Strigamia maritima*, *Isca Ixodes scapularis*, *Popi Phalangium opilio*, *Cscu Centruroides sculpturatus*, *Dsil Dysdera silvatica*, *Dpla Dolomedes plantarius*, *Lele Latrodectus elegans*, *Ptep Parasteatoda tepidariorum*, *Hgra Hylyphantes graminicola*, *Abru Argiope bruennichi*, *Tant Trichonephila antipodiana*, *Tcla Trichonephila clavata*.

|  | *Dmel* | *Tcas* | *Smar* | *Isca* | *Popi* | *Cscu* | *Dsil* | *Dpla* | *Lele* | *Ptep* | *Hgra* | *Abru* | *Tant* | *Tcla* |
| --- | --- | --- | --- | --- | --- | --- | --- | --- | --- | --- | --- | --- | --- | --- |
| Homeobox genes | 103 | 104 | 114 | 110 | 108 | 161 | 143 | 150 | 152 | 151 | 176 | 149 | 147 | 152 |
| Homeobox families | 83 | 85 | 90 | 89 | 91 | 90 | 79 | 87 | 89 | 91 | 86 | 88 | 87 | 89 |
| Families with paralogues from ancient TD(s) | 5 | 5 | 6 | 7 | 8 | 7 | 7 | 8 | 8 | 8 | 8 | 8 | 8 | 8 |
| Families with paralogues from recent TD(s) | 9 | 9 | 13 | 8 | 5 | 4 | 8 | 3 | 6 | 4 | 23 | 3 | 2 | 5 |
| Families with only one gene | 71 | 71 | 74 | 76 | 81 | 41 | 35 | 41 | 46 | 44 | 42 | 45 | 44 | 45 |
| Families with retained ohnologues | – | – | – | – | – | 46 | 41 | 44 | 41 | 44 | 37 | 40 | 40 | 40 |

In total, 90 described homeobox families were represented in our arthropod species set (Table 2), though some were absent in different species, such that no species had a gene from every family (Figure 2 A, Table S1). In addition, we also found several new homeobox genes. A few of these may represent new lineage-specific unnamed (“un”) families, temporarily called *ANTP-un1*, *PRD-un1*, *SINE-un1*, and *TALE-un1* based on the classes to which they likely belong. *ANTP-un1* is found only in the chelicerates, while *PRD-un1* and *SINE-un1* were found only in the araneids, and *TALE-un1* only in the *Trichonephila* spp. (Figure 2 A, Table S1). The ‘1’ is simply a placeholder to indicate that these genes have an orthologue in at least two species and to distinguish them from other unnamed or unclassified homeodomain sequences that have no similarity to described families and were found only in one species, sometimes in several copies.

**Figure 2.**
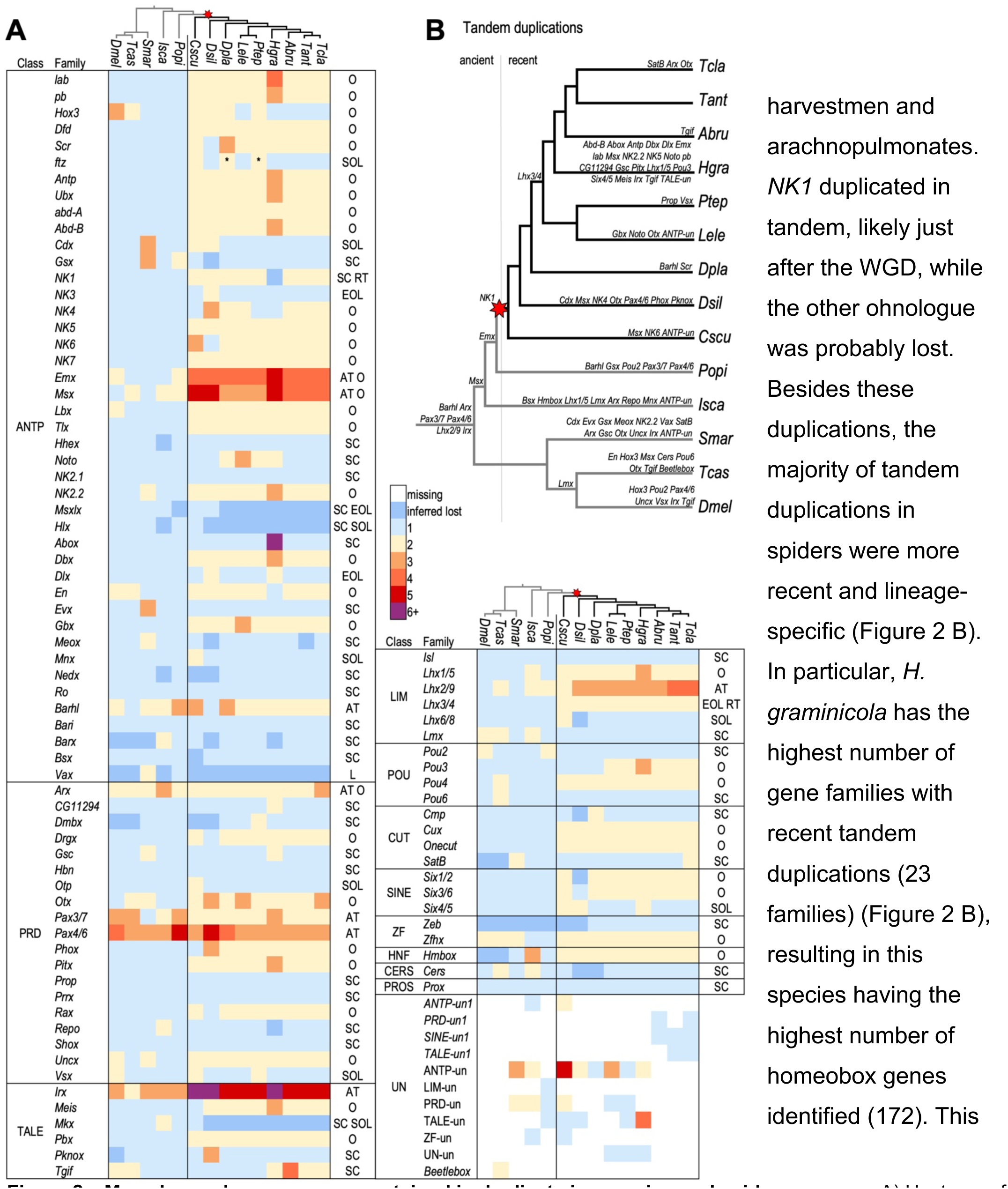
– Many homeobox genes were retained in duplicate in scorpion and spider genomes. A) Heatmap of homeobox genes by family. The relationships among duplicates (except for species-specific recent tandem duplications or family/species-specific losses) are indicated to the right. O: ohnologue retained; SOL: spider ohnologue lost; EOL: entelegyne ohnologue lost; AT: ancient tandem duplication; RT: recent tandem duplication; SC: single copy retained. *The second *ftz* ohnologues in *D. plantarius* and *P. tepidariorum* are likely pseudogenes as there are stop codons in the homeodomains. B) The inferred timing of ancient and recent tandem duplications among the included arthropod lineages. Gene families are listed above branches (as well as below for the root and *H. graminicola*). Species abbreviations are as in Table 2.

Comparing this taxonomic breadth of species in concert with genomic synteny enabled us to distinguish ancient and recent tandem duplicates from likely ohnologues and determine the timing of duplications relative to major radiations in arthropod evolution. We have defined ancient tandem duplicates as paralogues that are often linked or clustered in most spider genomes and date to a duplication node in an ancestor before the arachnopulmonate WGD, while recent tandem duplicates are also linked or clustered, but have occurred either after the WGD or in the outgroups. Six homeobox gene families, *Arx*, *Barhl*, *Irx*, *Pax3/7*, *Pax4/6*, and *Lhx2/9*, underwent ancient tandem duplication events that are shared by other arthropods and may have ancestry even deeper in the bilaterian phylogeny (Figure 2 B). Only three of these ancient tandem paralogues have been subsequently lost; just one *Lhx2/9* gene is found in *D. melanogaster*, and only one *Barhl* gene is found in *T. castaneum* and *D. silvatica*. Within the chelicerates, but predating the arachnopulmonate WGD, two further tandem duplications occurred. *Msx* likely duplicated in the common ancestor of ticks, harvestmen, and arachnopulmonates, and *Emx* likely duplicated in the ancestor of harvestmen and arachnopulmonates. *NK1* duplicated in tandem, likely just after the WGD, while the other ohnologue was probably lost.

Besides these duplications, the majority of tandem duplications in spiders were more recent and lineage-specific (Figure 2 B). In particular, *H. graminicola* has the highest number of gene families with recent tandem duplications (23 families) (Figure 2 B), resulting in this species having the highest number of homeobox genes identified (172). This could be an assembly artefact; however, overall the *H. graminicola* genome did not exhibit an exceptionally large number of duplicated genes (Zhu et al. 2022).

In contrast to tandem duplicates, dispersed paralogues consistently found on different chromosomes in surveyed spider genomes were inferred to be ohnologues. There were between 37 and 46 families with retained ohnologues in arachnopulmonates, and the families that were retained were largely consistent across species. All the Hox genes besides *Hox3* and *ftz* were retained as ohnologues, as previously reported for two spider species (*P. tepidariorum* and *Cupiennius salei*) (Schwager et al. 2017), as well as a few Super-Hox genes, namely *Dbx*, *En*, and *Gbx* (Figure 2 A). Many of the NK cluster genes were also retained as ohnologues, including *NK4*, *NK5*, *NK6*, *NK7*, *Lbx*, *Tlx*, as well as the ohnologues of ancient tandem paralogues of *Emx* and *Msx* (Figure 2 A).

Ohnologues were likely lost from several families following the WGD (Figure 2 A). Thirty-two families are found in single copies in scorpions and spiders (discounting subsequent lineage-specific tandem duplications of single-copy genes), indicating that an ohnologue was lost between the WGD and the divergence of scorpions and spiders. These include many of the Super-Hox families (*Abox*, *Dlx*, *Evx*, *Meox*, *Mnx*, *Nedx*, and *Ro*), the ParaHox family *Gsx*, most of the NK2 cluster families (*Hlx*, *Msxlx*, and *NK2.1*) and the HRO cluster family *Hbn*. Six families, including the Hox family *ftz*, the ParaHox family *Cdx* and the HRO cluster family *Otp*, were found in single copies in spiders, while both ohnologues were retained in the scorpion, indicating the loss of an ohnologue in a spider ancestor (Figure 2 A). Within spiders, an ohnologue was lost from four families in the entelegyne ancestor: *Dlx*, *Six4/5*, *Msxlx*, and *NK3*. *D. silvatica*, the sole synspermiatan representative, has lost the largest number of ohnologues from eighteen families, though the extent to which this applies across synspermiatans is not known. Only a few families underwent losses multiple times, namely *Hox3* in *C. sculpturatus*, *D. plantarius*, *Latrodectus elegans*, and the araneids; *Tgif* in the theridiids, *Dolomedes plantarius*, *D. silvatica* and *C. sculpturatus*; and *Barx* in both *H. graminicola* and *D. silvatica*. Only one family, *Vax*, was potentially lost before the WGD, or both ohnologues were lost in the arachnopulmonate ancestor (Figure 2 A). Though it was not reported previously (Leite et al. 2018), we identified a single *Vax* gene in the harvestman genome (Figure 2 A, Table S1).

### Homeobox gene clusters in spider genomes

#### The Hox cluster

Each of the spider genomes studied here contains two Hox clusters (Figure 3), consistent with previous descriptions for *P. tepidariorum* and other spiders (Schwager et al. 2007; Schwager et al. 2017; Fan et al. 2021; Sheffer et al. 2021; Wang et al. 2022). With the new *P. tepidariorum* chromosome-level assembly, we confirmed that each cluster is found on a different chromosome. Cluster B is typically intact, organised, and contains orthologues of all ten arthropod Hox genes, while cluster A often lacks *Hox3* and *ftz*, and is sometimes rearranged (Figure 3). There are some exceptions to this, however. For one, there have been several tandem duplications of *H. graminicola* Hox genes resulting in four *lab* paralogues, and three *pb*, *Antp*, *Ubx*, and *Abd-B* paralogues (Figure 3). While these duplications have occurred in both *H. graminicola* Hox clusters, cluster B is ordered, and contains the single orthologues of *Hox3* and *ftz*, while cluster A lacks those genes and has been rearranged (Figure 3). Furthermore, the Super-Hox gene *Dlx-A* is between *pb-A-2* and *Dfd-A* separated by 124 kb from *pb-A-2* and 380 kb from *Dlx-A* and *Dfd-A*. In *D. plantarius* there is a new gene, identified with highest similarity in the homeodomain to *Scr*, but not linked to either Hox cluster, possibly representing a divergent *Scr* paralogue (blue *‘S’* in Figure 3). In *D. silvatica*, both Hox clusters are rearranged, though one does contain both *Hox3* and *ftz*, while the other has lost only *ftz* (Figure 3). Pseudogenes of *ftz* were found in *D. plantarius* and *P. tepidariorum*, identified by the homeodomain disrupted by a stop codon. In *P. tepidariorum*, the *ftz*-pseudogene is not linked to either Hox cluster, while in *D. plantarius,* it is adjacent to *ftz*-B (Table S1, Figure S3 F).

**Figure 3.**
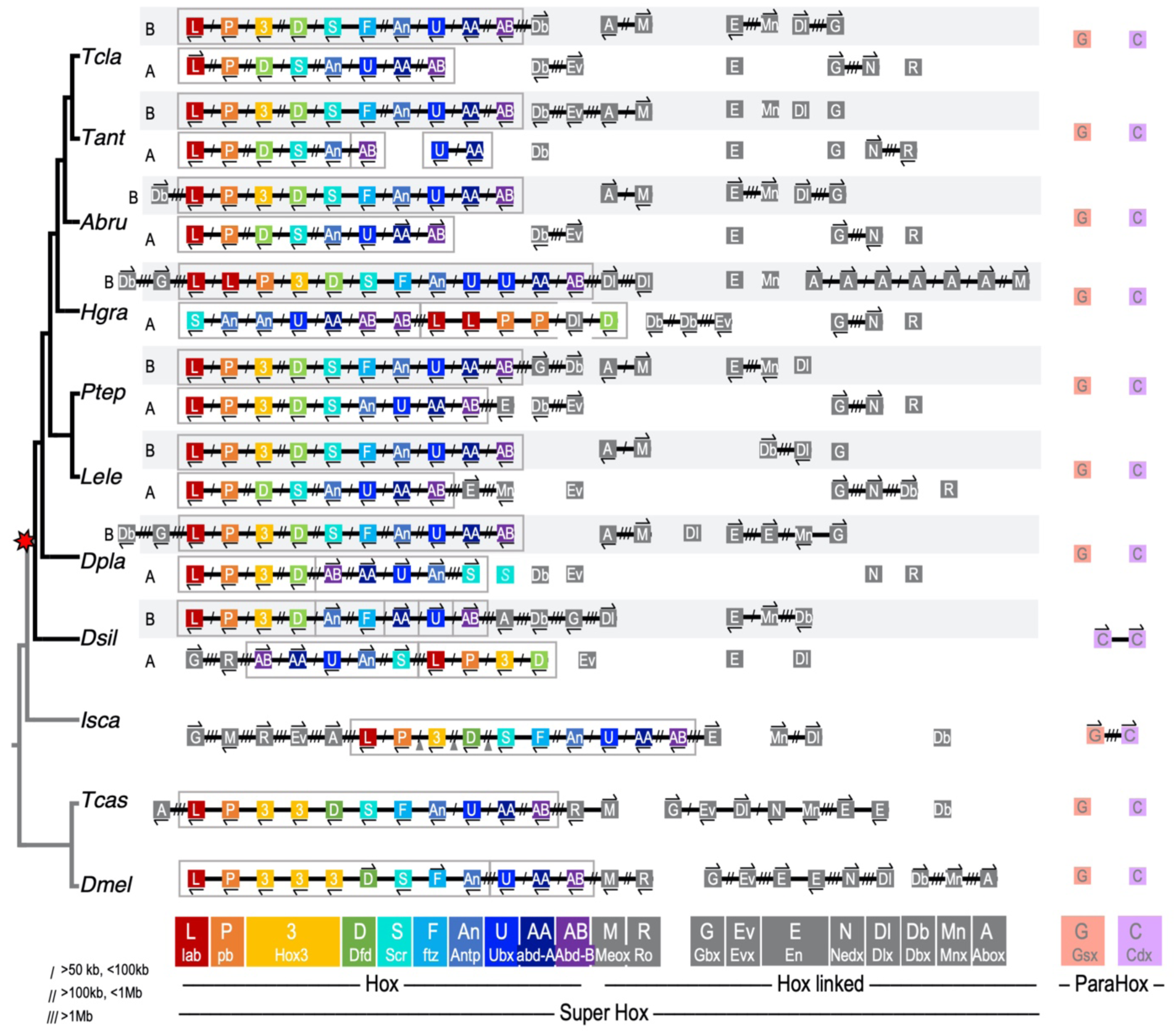
– Two ohnologous Hox clusters and a disintegrated arthropod ParaHox cluster are found in spiders. Genes are represented by coloured boxes; chromosomes are black lines. Gene orientation is indicated by arrows above or below genes, and distances are indicated by slashes between genes. Gene families are denoted by a one- or two-letter code, and by colour for Hox and ParaHox genes, or in grey for Hox-linked genes. ParaHox genes are coloured based on their relationship to Hox genes, *Gsx* with anterior genes and *Cdx* with posterior genes; *Xlox* was not found in any arthropod genome. Species abbreviations are as in Table 2.

The Super-Hox genes consist of the Hox cluster as well as several ANTP-class genes consistently found linked to each other and/or the Hox cluster across bilaterians (Table 1) (Butts et al. 2008; Ferrier 2016b). While many Super-Hox genes are still linked to one another and/or to the Hox cluster in the tick and insects, Super-Hox genes have dispersed in spider genomes and many Super-Hox families have also returned to the single copy (Figure 2 A, Figure 3). Amid the overall disintegration of the Super-Hox cluster, *Abox* and *Meox* are consistently found clustered in all the entelegyne species. Other remnants of linkage between Super-Hox genes include *Dbx* and *Gbx* linked to Hox cluster B in *H. graminicola*, *P. tepidariorum*, and *D. plantarius* (Figure 3). *En* is also often linked to *Mnx*, *Gbx* to *Nedx*, and *Dbx* to *Evx*, though these have more frequently been disrupted and dispersed, unlike the consistency of *Abox* and *Meox*. Synteny may be conserved by the constraint of regulatory elements or may result from the passive maintenance of ancestral linkage. While the Hox cluster has largely been retained in duplicate in spiders, most of the Super-Hox families have returned to the single copy and linkage has dispersed following the WGD.

The ParaHox cluster has largely disintegrated in spiders. Neither ParaHox gene is linked to the other, and both have returned to the single copy (Figure 3). The two *Cdx* paralogues in *D. silvatica* likely arose in a recent lineage-specific tandem duplication, as they form a two-gene cluster in only this species. In both insects, *Gsx* (*Ind*) and *Cdx* (*Cad*) are unlinked, while in the tick they are on the same chromosome but separated by a large distance (28 Mb) (Figure 3). This is consistent with the ancestral breakdown of the ParaHox cluster in an arthropod ancestor, and the further separation of the genes onto separate chromosomes in insects and spiders (Figure 3). It is not clear whether the retained ParaHox genes belong to the same ohnology group or not (i.e., both A or both B versus one A and one B), as there is only one ohnologue retained in any of the species surveyed.

#### The NK cluster

We found two ohnologous NK clusters in all spider genomes (Figure 4). One consists of *Msx1-A*, *NK4-A*, *NK3-A*, *Tlx-A*, *Lbx-A*, and *NK7-A* clustered on one chromosome, while *Msx1-B*, *Tlx-B*, *Lbx-B*, and *NK7-B* are found on a different chromosome, also with *NK3-B* and *NK4-B* in the species that have retained these genes (core NK clusters are delineated with grey boxes in Figure 4). Cluster B is less organised; *NK3-B* has likely been lost in entelegynes (estimated divergence 280-310 MYA) (Fernández, Kallal, et al. 2018; Shao and Li 2018) and *NK4-B* has likely been lost in *H. graminicola* and *P. tepidariorum*, independently. NK core cluster A is conserved across spiders, except that *Msx1-A* is not found in *L. elegans* or *P. tepidariorum*, which may indicate the loss of *Msx1-A* in a theridiid ancestor (est. 120 MYA) (Fernández, Kallal, et al. 2018). While NK cluster A is found in a conserved complement and order in almost all spider species analysed, it is rarely linked to any other genes belonging to the larger ancestral bilaterian NK complement (Table 1). In contrast, NK cluster B has a varying gene complement, having undergone lineage-specific gene losses and more rearrangements, making it more degenerate and less conserved between species. This second cluster is, however, often linked to other ancestral NK cluster genes *Hhex*, *Noto*, *Emx1-B*, *Emx2-B*, *NK5-B*, and *NK1*, which has undergone a tandem duplication, and to *NK6-B* and *Msx2-B* in all spiders (Figure 4). *NK6-A* is found linked to *Msx2-A* in all the entelegyne species, and *NK6* and *Msx2* genes can be found in cluster A and/or B in all spiders, suggesting this two-gene *NK6-Msx2* cluster predates the WGD and was retained in duplicate some species, while others have lost either *Msx2* or *NK6* ohnologues.

**Figure 4.**
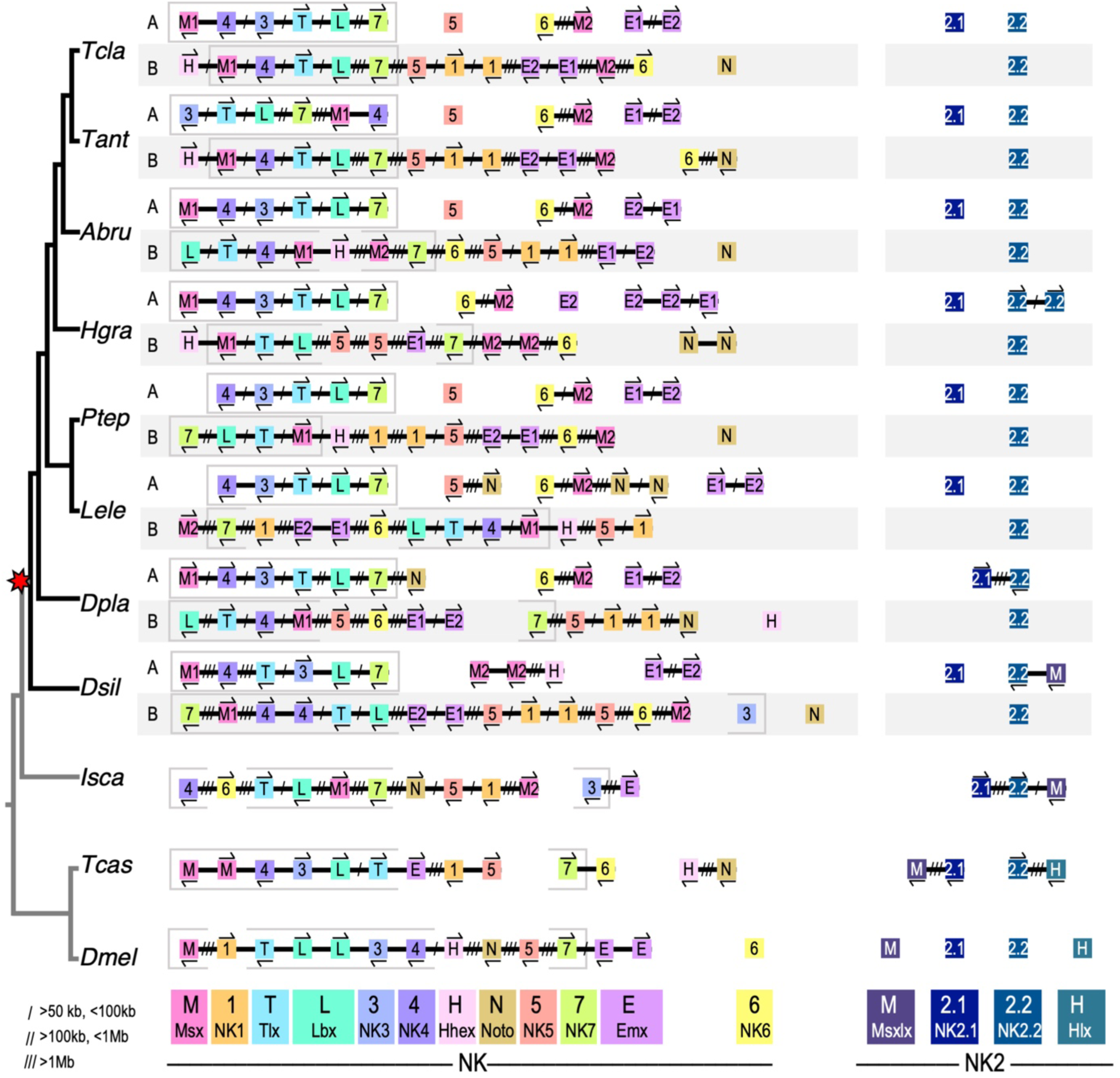
– Two ohnologous NK clusters are retained, and a disintegrated NK2 cluster is found in spiders. Genes are represented by coloured boxes; chromosomes are black lines. Gene orientation is indicated by arrows above or below genes, and distances are indicated by slashes between genes. Gene families are denoted by colour and a one- or two-letter code. The genes making up the spider core NK cluster are delineated by grey boxes. Species abbreviations are as in Table 2.

The NK2 cluster has largely broken down in spiders, with successive gene losses: first *Hlx* from chelicerates, and then *Msxlx* from entelegynes, though further lineage sampling may change this pattern (Figure 4). In all spiders, *NK2.2* is retained in duplicate on two separate chromosomes, indicating an ohnologous relationship, while *NK2.1* has returned to the single copy (Figure 4). *NK2.1* and *NK2.2* are unlinked in all spiders except for *D. plantarius*, indicating either that the cluster has separated twice, or alternatively, that different ohnologues have been lost in different lineages. Phylogenetics were not able to determine which ohnologue of *NK2.1* was retained in different species because no examined genome contains both ohnologues.

#### The SINE cluster

There are five SINE-class genes in two clusters in spiders, consistent with a duplication in the WGD followed by the loss of one *Six4/5* ohnologue. Spider SINE clusters likely underwent different rearrangements in different lineages (Figure 5). *D. sylvatica* has retained only one copy of each gene, while the entelegynes have retained B ohnologues of both *Six1/2* and *Six3/6* in a conserved cluster. In araneids, in cluster A all three genes are linked, though *Six4/5-A* is separated by a larger distance than that separating *Six1/2-A* from *Six3/6-A*, which remain tightly clustered. There has also been an inversion which may have occurred ancestrally among the theridiids that brought *Six4/5-A* closer to *Six3/6-A* than *Six1/2-A*. This indicates that the separation of *Six4/5-A* from the other two genes in araneids occurred after the divergence of the theridiids and that the three genes making up SINE cluster A were likely still tightly clustered in their shared ancestor. Among the outgroups, all three SINE genes are linked in the tick, and only two are linked in the fly, while none are linked in the beetle, indicating the spider SINE clusters have been retained more intact relative to other arthropod lineages.

**Figure 5.**
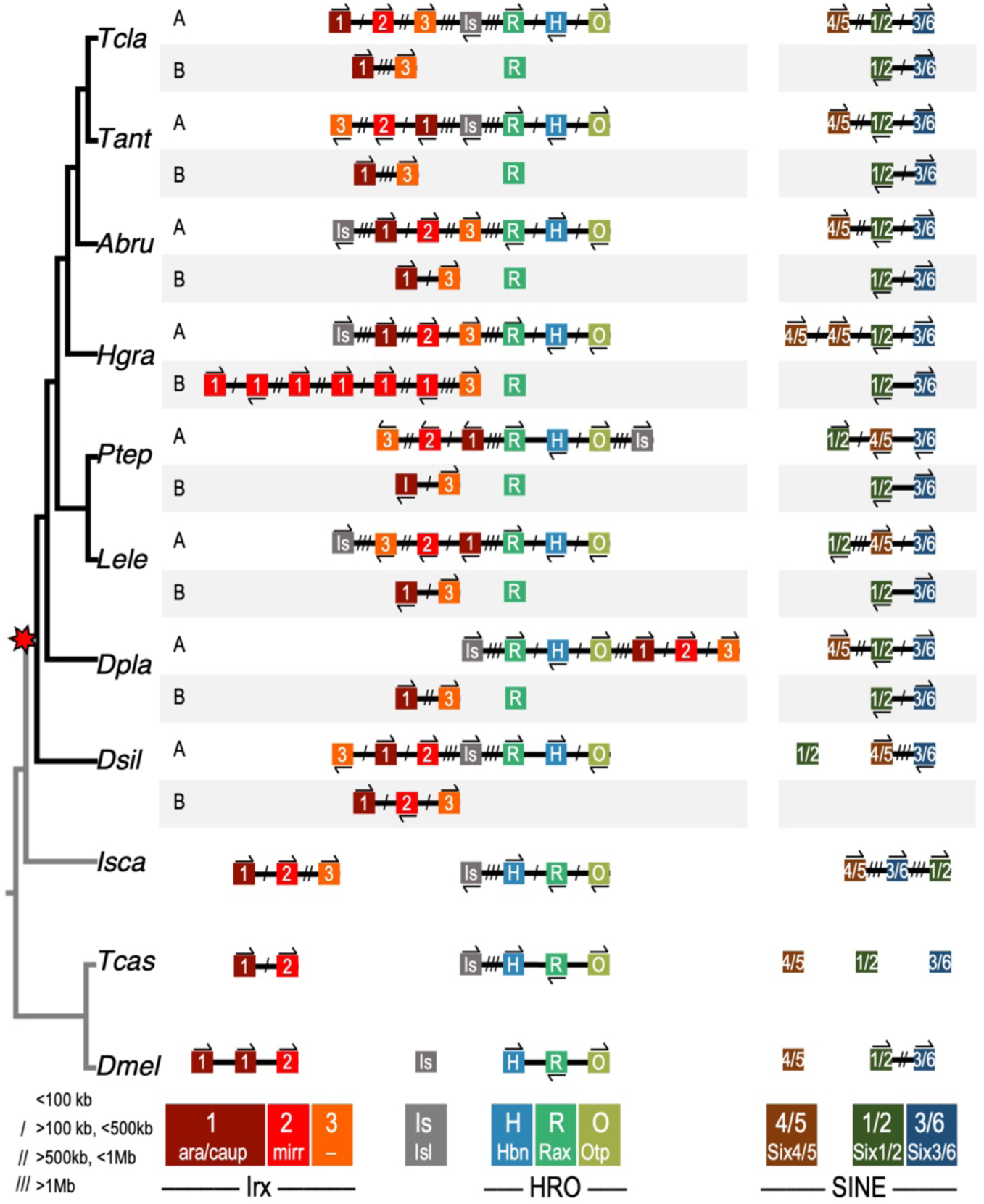
– Two ohnologous HRO clusters, one of which is linked to one of the two ohnologous *Irx* clusters, and two ohnologous SINE clusters are retained in spiders. Genes are represented by coloured boxes; chromosomes are black lines. Gene orientation is indicated by arrows above or below genes, and distances are indicated by slashes between genes. Gene families are denoted by colour and a one- or two-letter code. Species abbreviations are as in Table 2.

#### The HRO cluster

The PRD-class HRO cluster was so named because the order of the genes in the fly genome is *hbn* (*Hbn*), *Rx* (*Rax*), then *otp* (*Otp*); this order is conserved in insects and in the tick genome, indicating that it is likely the ancestral arthropod arrangement (Figure 5). In spiders, the HRO cluster is found intact, but reordered: *Rax*, *Hbn*, then *Otp* (Figure 5). There have also been several inversions of the orientations of the genes within the cluster, but the R-H-O order is conserved across spiders, suggesting this rearrangement occurred once ancestrally. It likely did not predate the WGD but occurred in a spider ancestor because the scorpion HRO cluster A order is *Hbn*, *Rax*, then *Otp*, as in insects and the tick (Table S1). As well as one conserved intact HRO cluster, spiders retained only *Rax* in duplicate, while there are two *Rax* and two *Otp* ohnologues retained in the scorpion (Figures 2 and 5).

The LIM-class gene *Isl* is consistently found linked to the HRO cluster (Figure 5). Ferrier (2016) suggested this pattern is evidence for the ancestral linkage between the LIM-class and PRD-class genes in the Giga-cluster of homeobox progenitor genes in an animal ancestor. The linkage of *Isl* to the HRO cluster has been observed across species with a most recent common ancestor in the ancestor of all animals. The consistent linkage pattern between these genes in spiders, the tick, and beetle genomes supports this association. This broad phylogenetic distribution makes this between-class linkage unlikely to be a convergent secondary association; rather it is more likely to be a conserved ancestral trait.

#### The Irx cluster

*Irx* genes have duplicated independently many times (see Table 1), giving rise to six genes in spiders. Each of the ohnologous *Irx* clusters contains one of each gene type, *Irx1*, *Irx2*, and *Irx3*, though *Irx2-B* has been lost from one of the clusters in the entelegynes (Figure 5). Consistent with the findings of Kerner et al. (2009), insect *Irx* genes fall into two clades, one containing orthologues of *D. melanogaster mirr* and the other with orthologues of *ara* and *caup* (Table 1 and Figure S1). Spider and scorpion *Irx1-A* and *Irx1-B* group with *ara* and *caup*, while spider and scorpion *Irx2-A* and *Irx2-B* group with *mirr* (Figure S1). The exact relationships between *Irx3* and other arthropod genes remains somewhat unclear. There are three *Irx* genes in the centipede *S. maritima* and the millipede *Trigoniulus corallinus* genomes (Chipman et al. 2014; So et al. 2022), one of which is not contained within either the *Irx1/ara/caup* or *Irx2/mirr* clade, and is resolved as sister to the *Irx3* clade in the *Irx* gene phylogeny, albeit with very low bootstrap support (Figure S1).

The potential *Irx3* is clustered with *Irx1* and *Irx2* in the *S. maritima* genome in the same relative position as spider *Irx3* (Figures 5 and S3 C). There is a clear orthologue of each of the three arachnopulmonate *Irx* gene families in both the harvestman and tick genomes, indicating the three gene types originated in or before the chelicerate ancestor, and possibly *Irx3* genes were lost from an insect ancestor after the divergence of Myriapoda (est. 520 MYA) (Fernández, Edgecombe, et al. 2018). According to Kerner et al. (2009), *Irx* genes duplicated independently in spiralians, ecdysozoans, and deuterostomes, suggesting that all paralogues arose from a single ancestral bilaterian *Irx* gene. From our phylogeny (Figure S1), we can tentatively infer that the duplication that gave rise to fly paralogues is ancestral to the arthropods, and that there was a three-gene cluster in the ancestral chelicerate that further duplicated in the arachnopulmonate WGD resulting in up to six spider *Irx* paralogues in two ohnologous clusters.

In spiders, one of the two ohnologous *Irx* clusters is also linked to the HRO+*Isl* cluster (Figure 5). Since this association occurs only to *Irx* cluster A, while *Irx* cluster B is consistently not linked to the second *Rax* paralogue (Figure 5, Figure S3 E-L), a likely explanation is that one *Irx* cluster translocated to be linked to HRO+*Isl* following the WGD. We infer it occurred secondarily, i.e., on only one ancestral arachnopulmonate chromosome, rather than deeper in animal ancestry due to the consistent lack of linkage between HRO+*Isl* with *Irx* genes in any other species examined here, in contrast to the linkage between HRO and *Isl* found across bilaterians.

### Conserved macrosynteny of spider genomes

To contextualise our observations of homeobox cluster conservation and rearrangements, we explored macrosynteny across spider chromosomes. We used *H. graminicola* as the reference because the *D. silvatica* genome is highly rearranged compared to all entelegynes in this study, and *H. graminicola* represents an early-branching lineage realative to the majority of other entelegynes (Figure 6, Figure S4 A). Across entelegynes, we detected clear patterns of orthology between chromosomes, and consistency in the synteny blocks in which certain homeobox clusters are found, but these patterns were not as evident when compared to the sole synspermiatan representative *D. silvatica* (Figure 6, Figure S4 A). Depictions of locations of all homeobox genes across genomes can be found in Figure S3.

**Figure 6.**
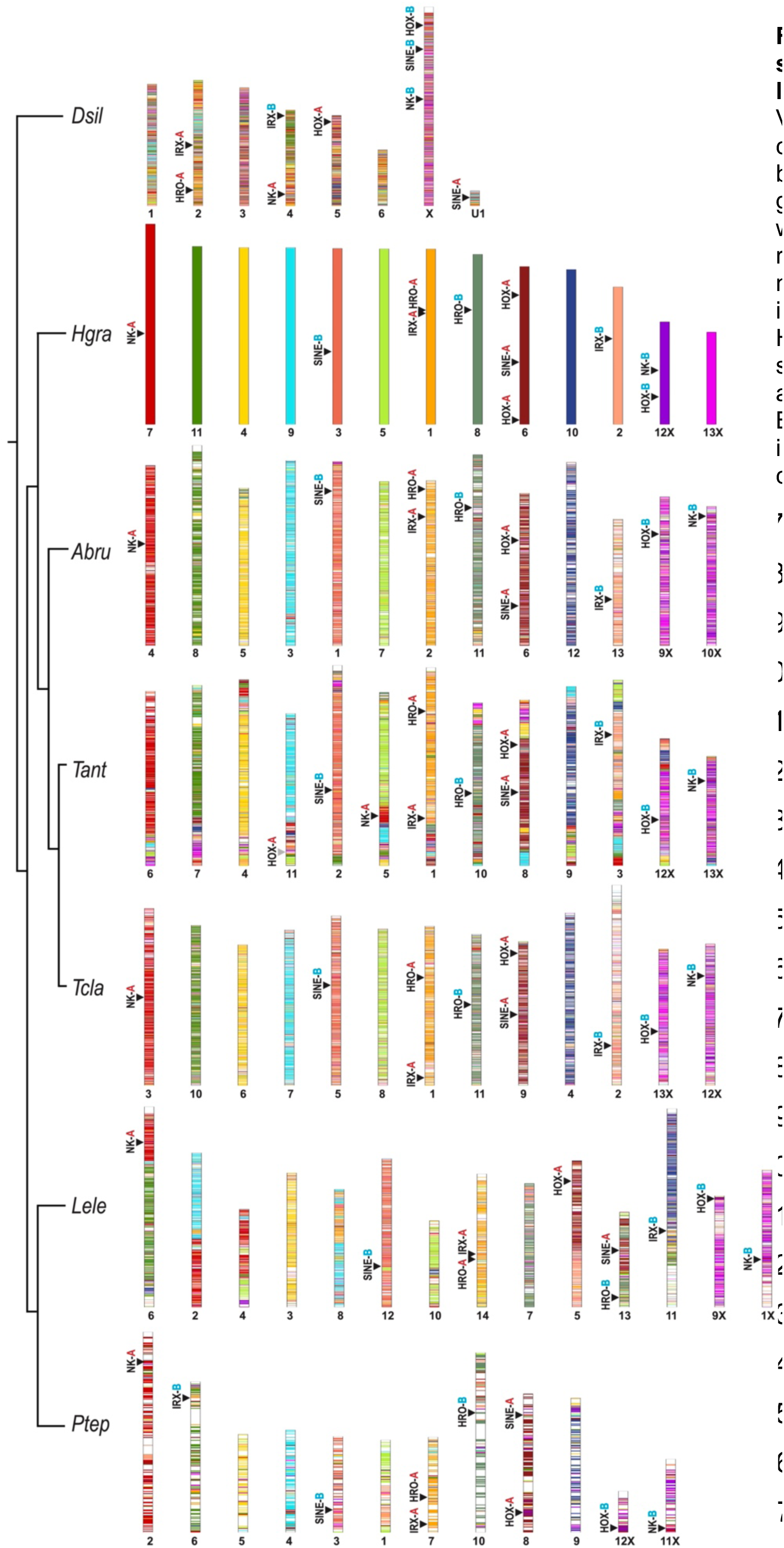
– Conserved macrosynteny of spider chromosomes and associated locations of homeobox gene clusters. Vertical bars show stacks of genes on each chromosome representing a macrosynteny block, with *H. graminicola* as the reference genome. Chromosomes of other species were ordered to show the orthologous relationships to *H. graminicola*. Chromosome numbers are denoted below, with ‘X’ indicating sex chromosomes. Locations of HOX, NK, IRX, SINE and HRO clusters are shown with black arrows, along with the assignment for cluster A (red) versus cluster B (blue). The grey triangle in *T. antipodiana* indicates the partial translocation of Hox cluster A.

In each araneid analysed here, there are 13 chromosomes; however, in the theridiids, there are 12 in *P. tepidariorum* and 14 in *L. elegans* (Figure 6, Figure S4 A). In *P. tepidariorum*, the synteny block corresponding to chromosome 2 in *H. graminicola* (coral pink in Figure 6) appears to have been mostly lost or distributed throughout the genome, and *Irx* cluster B which belongs to this block has been relocated (Figure 6, Figure S4 A). In *L. elegans* there appear to have been large fission and fusion events on chromosomes 2, 4, 5, 6, 8, 11, and 13. Such events in *L. elegans* are consistent with Hox cluster A and SINE cluster A being found on separate chromosomes, as well as the rearrangement of *Irx* cluster B (Figures 6 and S3 G, Figure S4 A).

Sex chromosomes have been identified in several spider species (Figure 6 and Figure S4 A) (Sheffer et al. 2021; Zhu et al. 2022; Miller et al. 2023). Interestingly, Hox cluster B and NK cluster B are both found on the same sex chromosome in *H. graminicola* (12) and *D. silvatica* (X), but on separate sex chromosomes in other entelegynes, whereas Hox cluster A is located on an autosome (Figure 6, Figure S4 A). Across entelegynes, NK cluster A is found in the same conserved synteny block corresponding to a full chromosome in all species except for *Trichonephila antipodiana*. In this species, a rearrangement brought NK cluster A onto another chromosome, and this is the only entelegyne species in which the NK cluster A is rearranged (Figure 4). Overall, this analysis illustrates the conservation of macrosynteny blocks and chromosomes across entelegyne genomes and shows several chromosomal rearrangements that have affected homeobox gene clusters.

### Analysis of selection on NK cluster genes

We next sought to understand how homeobox genes have evolved following WGD and tandem duplications. In the NK cluster, we identified families with single-copy orthologues, ancient tandem paralogues, and WGD ohnologues. To determine if genes with different relationships were under different selective pressures, we compared the aBSREL estimated ω values calculated across coding sequence phylogenies for seven NK cluster families with different inferred duplication histories: *Msx*, *NK4*, *NK3*, *Tlx*, *Lbx*, *NK7*, and *Emx*. While none of the focal ohnologues or tandem paralogues were found to be under positive selection, certain genes from the pre-duplicate outgroups were. *NK7* was found to be under positive selection in *D. melanogaster* and *I. scapularis*, *Tlx* also in *I. scapularis*, and *NK3* in *T. castaneum*. However, certain ohnologues did have longer branches separating them from other ohnologues or the single-copy outgroup genes in the aBSREL-generated phylogenies, indicating a higher substitution rate. These include *Msx1* in arachnopulmonates and the harvestman, *NK3-A* in the entelegynes, and *Lbx-B* in the scorpion. No consistent pattern was detected between A versus B ohnologues, despite NK cluster A being more organised, nor was there a clear pattern between ohnologues versus tandem duplicates. Supplementary material underpinning these results is available on GitHub (see Materials and Methods).

In the one-to-one comparisons between *P. tepidariorum* and *P. opilio* genes, and between *P. tepidariorum* ohnologues, we calculated dN/dS ratios with codeML to detect positive selection. For all NK cluster genes, none approached 1; all were very low, ranging from 0.0011 (between *P. opilio Msx2* and *P. tepidariorum Msx2-A*) to 0.1423 (between *P. tepidariorum Emx1-A* and *Emx1-B*) (Tables 3 and S3). There are some patterns among gene families, e.g., lower dN/dS ratios for *Tlx* ohnologues compared to a consistently higher rate for *NK7* ohnologues. The *Emx1* and *Emx2* and *Msx1* and *Msx2* ohnologues had consistently low dN/dS values when compared to their *P. opilio* orthologues but varied when compared between ohnologues in *P. tepidariorum*. Overall, these low dN/dS ratios likely reflect the highly conserved nature of homeobox genes.

**Table 3.** – The ratio of non-synonymous to synonymous substitution rates between *P. tepidariorum* and *P. opilio* NK cluster genes shows no evidence for positive selection among paralogues. Pairwise dN/dS values were calculated with codeML for each *P. opilio* and *P. tepidariorum* NK cluster gene. The bars at the right show the dN/dS value relative to the black bar (length of 1) at the top. The relationship between each gene pair is indicated in the third column, where 1 to 1 indicates direct orthology (the loss of an ohnologue), 1 to 2 indicates the retention of two ohnologues in the spider, O is the comparison between the two spider ohnologues, and T is the comparison between tandem paralogues. For families with ancient tandem duplications (*Msx* and *Emx*), gene names have been updated to reflect the inferred relationships between paralogues in *P. opilio* and subsequent ohnologues in *P. tepidariorum*, such that, e.g., *P. opilio Emx1* is the pro-orthologue of *P. tepidariorum Emx1-A* and *Emx1-B*, and the same for *P. opilio Emx2* and *P. tepidariorum Emx2-A* and *Emx2-B*. Raw substitutions (S and N) and substitution rates (dS and dN) as calculated with codeML can be found in Table S3.

| *Popi* | *Ptep* |  | dN/dS |
| --- | --- | --- | --- |
| NK4 | NK4 | 1 to 1 | 0.0673 |
| NK3 | NK3 | 1 to 1 | 0.0584 |
| *Tlx* | *Tlx-B* | 1 to 2 | 0.0060 |
|  | *Tlx-A* | 1 to 2 | 0.0094 |
|  | *Tlx-A* x *Tlx-B* | O | 0.0042 |
| *Lbx* | *Lbx-B* | 1 to 2 | 0.0421 |
|  | *Lbx-A* | 1 to 2 | 0.0299 |
|  | *Lbx-A* x *Lbx-B* | O | 0.0076 |
| *NK7* | *NK7-B* | 1 to 2 | 0.0642 |
|  | *NK7-A* | 1 to 2 | 0.0430 |
|  | *NK7-A* x *NK7-B* | O | 0.0575 |
| *Msx1* | *Msx1-B* | 1 to 1 | 0.0104 |
| *Msx2* | *Msx2-A* | 1 to 2 | 0.0011 |
|  | *Msx2-B* | 1 to 2 | 0.0106 |
|  | *Msx2-A* x *Msx2-B* | O | 0.0051 |
|  | *Msx1-B* x *Msx2-B* | T | 0.0845 |
| *Emx1* | *Emx1-A* | 1 to 2 | 0.0123 |
|  | *Emx1-B* | 1 to 2 | 0.0128 |
| *Emx2* | *Emx2-A* | 1 to 2 | 0.0120 |
|  | *Emx2-B* | 1 to 2 | 0.0094 |
|  | *Emx1-A* x *Emx1-B* | O | 0.1423 |
|  | *Emx2-A* x *Emx2-B* | O | 0.0118 |
|  | *Emx1-A* x *Emx2-A* | T | 0.0188 |
|  | *Emx1-B* x *Emx2-B* | T | 0.0124 |

### Developmental expression of NK cluster genes in P. tepidariorum and P. opilio

It was previously shown that several ohnologues of spider homeobox genes had undergone sub- and/or neofunctionalisation with respect to their single-copy orthologues in the harvestman (Leite et al. 2018). To better understand the roles of homeobox genes following WGD in the arachnopulmonate ancestor, we assayed the expression of NK cluster genes in the spider *P. tepidariorum* in comparison to the harvestman *P. opilio*.

#### Single-copy genes

Only one copy each of *NK3* and *NK4* were found in both *P. opilio* and *P. tepidariorum*, indicating the loss of an ohnologue of each from the spider following the WGD. The single spider *NK3* is strongly expressed in the developing heart and is weakly expressed around the stomodaeum and in the head (Figure 7 A-E). This is very similar to the single spider *NK4*, which is also expressed strongly in the heart (Figure 7 F-J).

**Figure 7.**
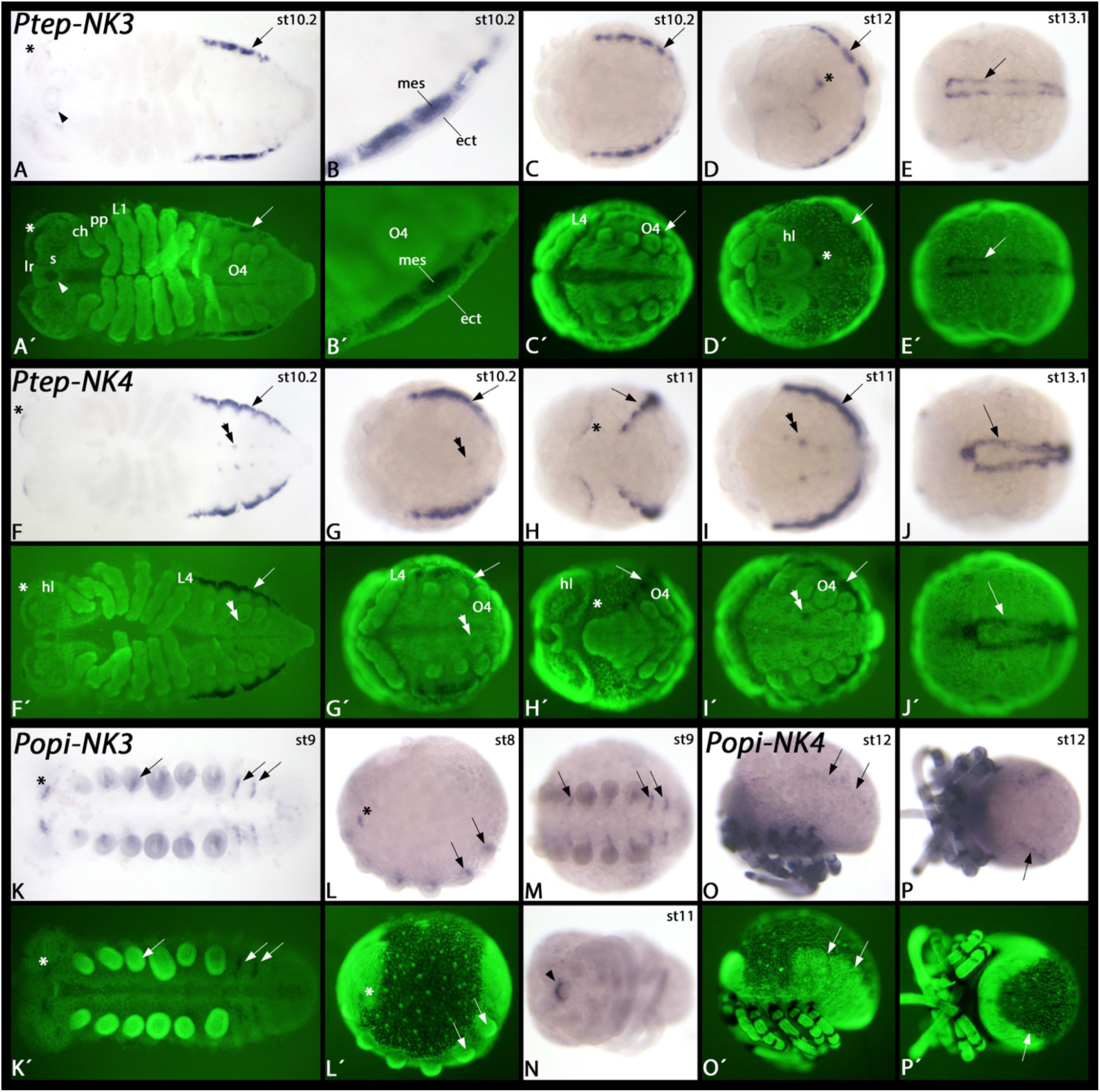
– Expression of *P. tepidariorum* and *P. opilio NK3* and *NK4*. In all panels, anterior is to the left. Panels A-C, F, G, I, K, M, and N show ventral views. Panels D and H show anterior-dorsal views. Panels E, and J show dorsal views. Panels L and O show lateral views and dorsal up. Panel P shows a posterior-dorsal view. Embryos in panels A, F and K are flat-mounted. Panel B shows a magnification of the posterior region of a flat-mounted embryo. Panels Á-Ĺ, Ó and P′ show SYBR green stained embryos as shown in panels A-L, O and P. Developmental stages of *P. tepidariorum* (after Mittmann and Wolff (2012)) and *P. opilio* (after Gainett et al. (2022)) are indicated in the upper right corner of each panel with a bright field image. Arrows in panels A, C-J, O and P mark expression of *NK3* and *NK4* in the developing heart of *P. tepidariorum*, and expression of *NK4* in the developing heart of *P. opilio*. The arrowhead in panel A points to expression in the labrum. The asterisk in panels A and D mark expression in the head. In panel B, expression in the mesoderm is marked (note absence of expression in the overlying ectoderm). Double arrowheads in panels F, G and I point to dot-like expression of *NK4* in the opisthosoma of the spider. Asterisks in panels F and H mark stripes of expression in the anterior of the head. Asterisks in panels K and L mark two domains of *NK3* expression in the head of the harvestman. Arrows in panels K-M mark short segmental stripes of *NK3* expression. The arrowhead in panel N marks expression in the labrum at late developmental stages. Abbreviations: ch, chelicera; ect, ectoderm; hl, head lobe; L1-L4, first to fourth walking leg; lr, labrum; mes, mesoderm; O, opisthosomal segment; pp, pedipalp; s, stomodaeum.

Unlike *NK3*, *NK4* is also expressed in a dot-like fashion in the ventral nervous system of the opisthosoma, and at the anterior rim of the head lobes (Figure 7 asterisks and double arrowheads in Figure 7 F-I). We did not detect any *NK3* expression in the developing heart of the harvestman; instead *NK3* is expressed in a segmental fashion ventral to the base of the appendages and in similar short transverse segmental stripes in the opisthosomal segments (Figure 7 K-N). The expression of harvestman *NK3* is thus more like the expression of its orthologue in *D. melanogaster* (Azpiazu and Frasch 1993) and the millipede *Glomeris marginata* (Figure S5) than that of the spider. Expression of *NK4* in the harvestman is restricted to the developing dorsal vessel at late developmental stages (Figure 7 O-P), a pattern that is conserved among panarthropods (Azpiazu and Frasch 1993; Janssen and Damen 2008; Treffkorn et al. 2018).

#### Ancient tandem paralogues

Both *Emx* and *Msx* families have likely undergone an ancient tandem duplication, resulting in two paralogues of each gene in the harvestman genome, with four *Emx* and three *Msx* paralogues retained in *P. tepidariorum*. In previous transcriptome-based analyses of *P. opilio*, the highly similar sequences for each *Emx* paralogue were merged into a single transcript model, and only a partial sequence was found for *Msx1*, split across two different transcript models (Sharma et al. 2012), but the genome revealed distinct loci for each paralogue. In *P. tepidariorum*, Leite et al. (2018) described *Emx1-A* and *Emx2-A* (previously designated *Emx4* and *Emx3*, respectively) expression in the precheliceral region and in patches in each segment along the ventral midline, and *Emx1-B* and *Emx2-B* expression (previously designated *Emx2* and *Emx1*) in the anterior of each opisthosomal segment, with *Emx1-B* also expressed at the base of the prosomal appendages. *P. opilio Emx2* (previously designated *Emx*) was detected at the base of the appendages and in patches along the ventral midline in stage 10-11 embryos (Leite et al. 2018).

Very similar to its paralogue *Emx2*, *P. opilio Emx1* is expressed in all posterior segments including the cheliceral segment (Figure 8). The sharp anterior border of expression is conserved among other arthropods, including spiders (Walldorf and Gehring 1992; Simonnet et al. 2006; Schinko et al. 2008; Birkan et al. 2011; Janssen 2017; Leite et al. 2018) and an onychophoran (Janssen 2017). Indeed, the overall expression pattern of *P. opilio Emx1* is identical with that of the previously reported *Emx2*, with the exception that expression of *Emx1* is detected earlier.

**Figure 8.**
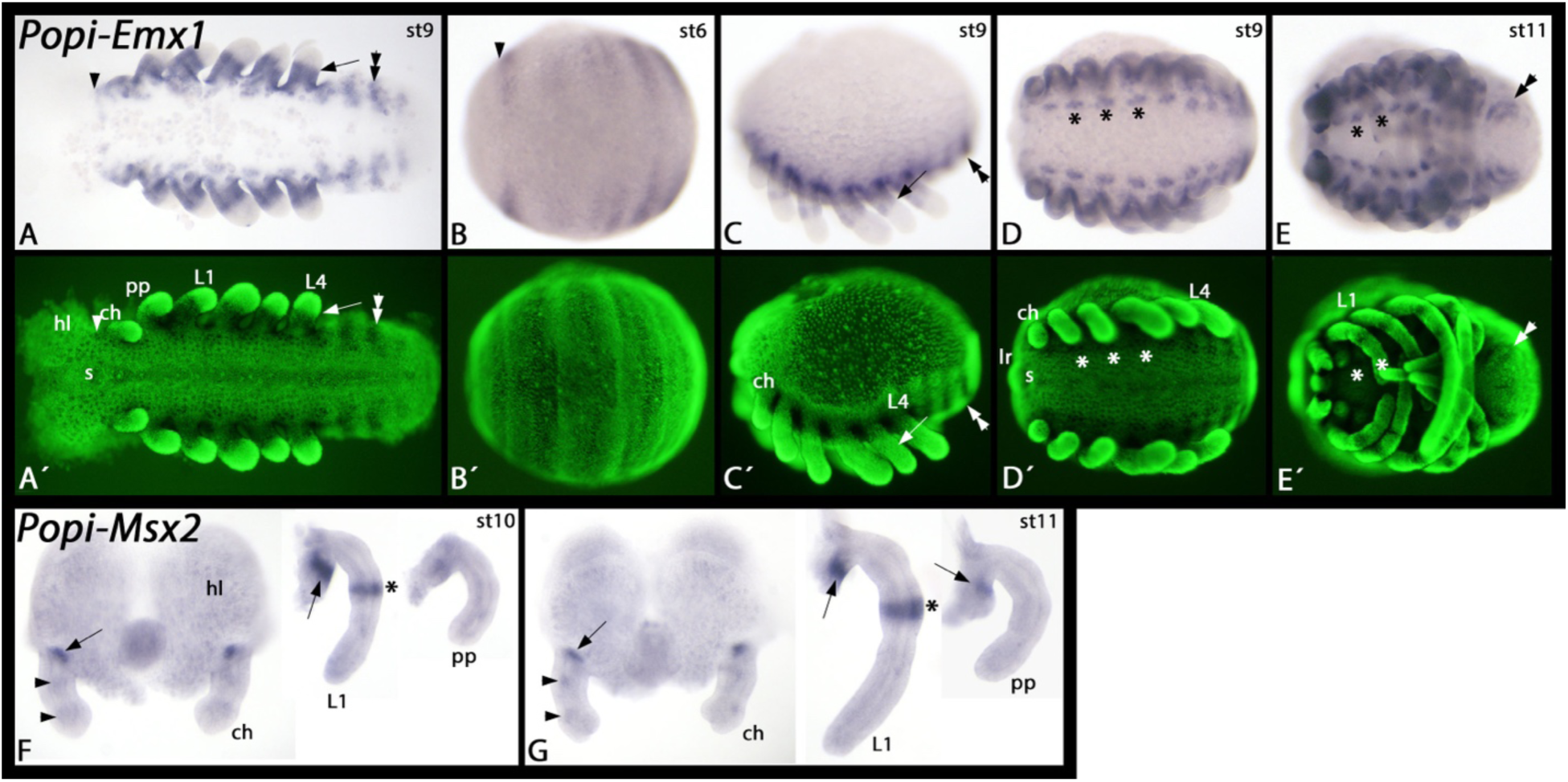
– Expression of *P. opilio Emx1* and *Msx2*. In panels A-E, anterior is to the left and ventral views (except panel C, lateral view and dorsal up). Panels F and G each show dissected head lobes and chelicerae, a walking leg and a pedipalp. The dissected heads are shown with anterior up, ventral views. The dissected legs and pedipalps present lateral views, dorsal to the right. Panels Á-É show SYBR green stained embryos as shown in panels A-E. Developmental stages after Gainett et al. (2022) are indicated in the upper right corner of each panel with a bright field image. Arrowheads in panels A and B mark the sharp anterior border of *Emx1* expression anterior to the cheliceral segment. The arrows in panels A and C mark sharp distal border of expression in the appendages. Asterisks in panel D mark expression in the ventral nervous system, ventral to the base of the appendages. Double arrowheads in panels A, C and E mark expression in the opisthosoma. Arrows in panels F and G point to expression at the base of the appendages. Arrowheads in panels F and G point to dot-like expression in the chelicerae. Asterisks in panels F and G mark a ring-like expression in the legs. Abbreviations: ch, chelicera; hl, head lobe; L1-L4, first to fourth walking leg; lr, labrum; pp, pedipalp.

According to Leite et al. (2018), *P. tepidariorum Msx1-B* (previously designated *Msx3*) is expressed in the base of the prosomal appendages at stage 11, while *Msx2-B* (previously designated *Msx2*) is expressed in the chelicerae at stage 12, and *Msx2-A* (previously designated *Msx1*) is expressed in each segment along the ventral midline in stages 8 and 10 (Akiyama-Oda and Oda 2020). In *P. opilio*, *Msx1* (previously designated *Msx*) was detected in a stripe in each segment at stage 7 and along the ventral midline in each segment at stage 13 (Leite et al. 2018). In contrast, we detected *P. opilio Msx2* expression only at late developmental stages and in a very restricted dot-like patch at the base of the appendages, and a ring-like domain in the legs (Figure 8 F-G). Thus, *P. opilio Msx1* expression in the segments and along the ventral midline is similar to *P. tepidariorum Msx2-A* expression, while *P. opilio Msx2* expression at the base of appendages is similar to *P. tepidariorum Msx1-B* expression. *P. opilio Msx2* expression was also detected in a small region of the chelicerae, matching the expression of *Msx2-B* in stage 12 *P. tepidariorum* that was observed by Leite et al. (2018) and interpreted as a novel expression domain. Our description of a matching expression pattern from a previously unknown harvestman gene, *Msx2*, means that the inferred neofunctionalisation is more likely a case of subfunctionalisation between the tandem *Msx* paralogues.

#### Ohnologues

Three other NK families, *NK7*, *Tlx*, and *Lbx*, are found in the single copy in *P. opilio* and have two retained ohnologues in *P. tepidariorum* (Figure 4). The two ohnologues of *P. tepidariorum Tlx* are both expressed in the distal region of the legs, but in complementary patterns (Figure 9 A-M). *Tlx-A* is first expressed in the complete distal region of the appendages and restricts into two rings of stronger expression around stage 10 (Figure 9 G-H). *Tlx-B* expression only begins after this point and is strongest in the region between the two rings of *Tlx-A* expression (Figure 9 M). Apart from its expression in the developing appendages, *Tlx-A* is also expressed in the dorsal tissue of the opisthosoma in a pattern that suggests a role in heart development (Figure 9 B, D-E), and at later stages, in the ventral opisthosoma (Figure 9 F). In contrast, *Tlx-B* is only expressed in the opisthosoma at the very posterior at late developmental stages, after segment addition is complete (Figure 9 J). Unlike *Tlx-A*, *Tlx-B* is expressed in the form of a transverse stripe in the developing head at late developmental stages (double arrowheads in Figure 9 K) at the anterior edge of non-neurogenic ectoderm that overgrows the neurogenic ectoderm of the head lobe from the anterior.

**Figure 9.**
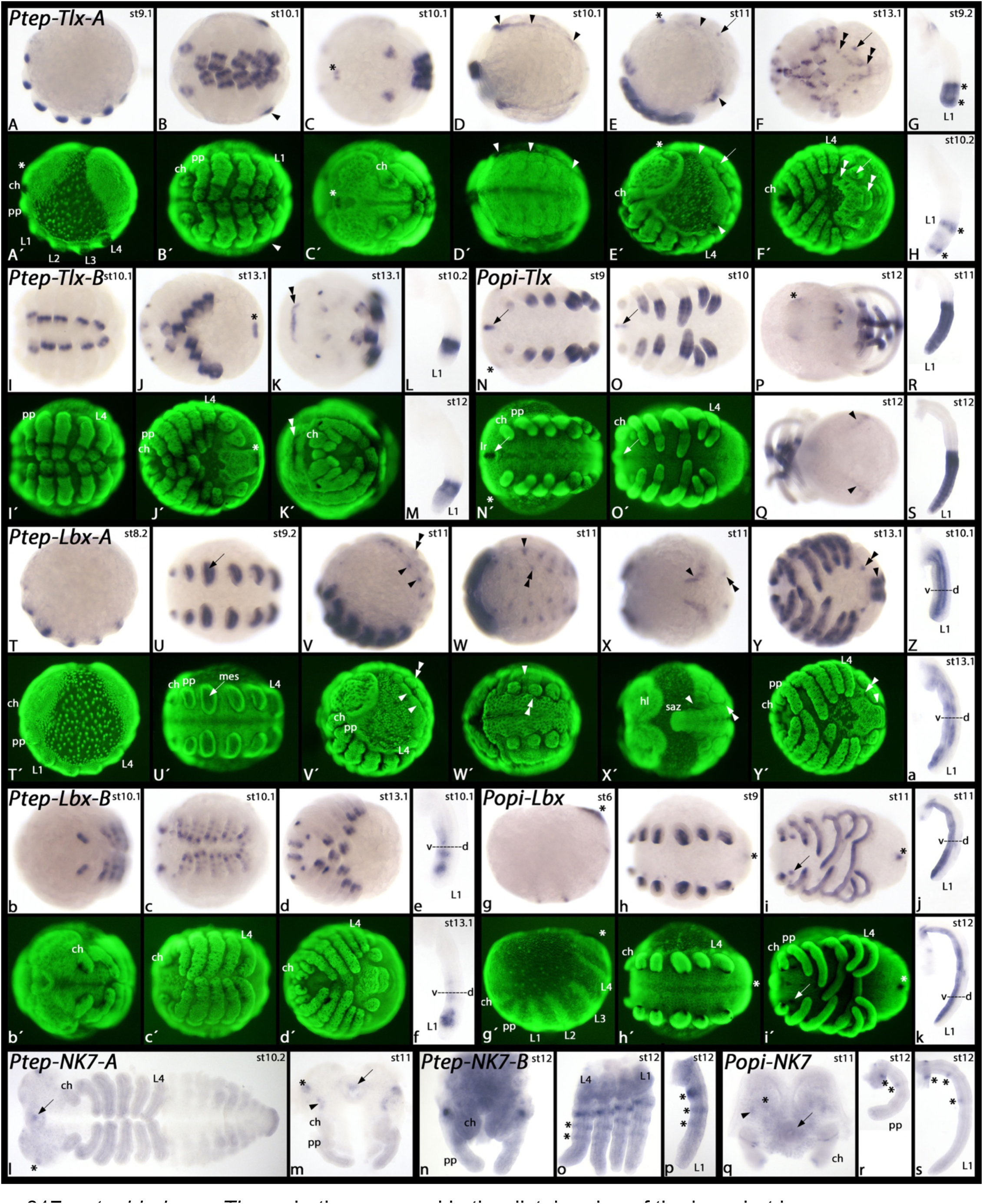
– Expression of *P. tepidariorum* and *P. opilio Tlx*, *Lbx* and *NK7* ohnologues. In all panels showing whole embryos, anterior is to the left. Dissected heads in panels m, n, and q, are anterior up. Dissected appendages are shown by lateral views, dorsal to the right. Panels A, E, T, V, and g show lateral views and dorsal up. Panels B, C, D, F, I, J, N, O, U, W, Y, b, c, d, h, and i show ventral views. Panels K and P show anterior views. Panel Q shows a posterior view. Panel X shows a dorsal view. The embryo in panel I is flat-mounted. Panels Á-F, I-K′, Ń, Ó, T′-Ý, b′-d′, and ǵ-í show SYBR green stained embryos as shown in panels with corresponding bright field panels. Developmental stages of *P. tepidariorum* (after Mittmann and Wolff (2012)) and *P. opilio* (after Gainett et al. (2022) are indicated in the upper right corner of each panel with a bright field image. Arrowheads in panels B, D, and E mark dorsal expression in the opisthosoma. The asterisks in panels C and E mark expression in the labrum. The arrows in panels E and F mark expression in the posterior spinneret, and the double arrowhead marks ventral expression in the opisthosoma. Asterisks in panels G and H mark two stripes of stronger expression within the distal region of *Tlx-A* expression in the legs. The asterisk in panel J marks expression of *Tlx-B* in the very posterior of the embryo. The double arrowhead in panel K marks a transverse stripe of expression in the head. The arrows in panels N and O mark expression of *Tlx* in the labrum of the harvestman. The asterisk in panel P marks expression in the head lobes. The arrow in panel U points to expression in the mesoderm of the leg. Arrowheads in panels V-Y point to dorsal expression in the opisthosoma and double arrowheads point to expression ventral to the base of the opisthosomal appendages. The asterisks in panels g-i mark expression in the very posterior of the embryos. The arrows in panels l, m, and q point to expression in the labrum. Asterisks and arrowheads in panels m and q mark expression of *NK7* in the head of the spider and the harvestman. Asterisks in panels o, p, r, and s mark expression in the appendages. Abbreviations: ch, chelicera; L1-L4, first to fourth walking leg; mes, mesoderm; pp, pedipalp; saz, segment addition zone.

The single *Tlx* gene of the harvestman shares many aspects with one or both spider ohnologues, indicating the spider *Tlx* ohnologues have subfunctionalised. *P. opilio Tlx* is expressed in the distal half of the developing appendages (Figure 9 R-S), reflecting the combined expression of both spider *Tlx* ohnologues (Figure 9 G-H, L-M). Like *P. tepidariorum Tlx-A*, *P. opilio Tlx* is expressed in dorsal tissue of the opisthosoma and in the labrum (Figure 9 N-O), and like *P. tepidariorum Tlx-B*, *P. opilio Tlx* is expressed at the anterior edges of the head lobes (asterisk in Figure 9 P).

Expression of *P. tepidariorum Lbx-A* first appears in the mesoderm of the outgrowing prosomal limb buds (Figure 9 T-U). Later, expression in the appendages is also seen in the ventral ectoderm, but the dorsal ectoderm remains free of expression (Figure 9 W-Y). Additional expression of *Lbx-A* is found in the dorsal ectoderm of the opisthosoma and in segmental patches in the opisthosoma (Figure 9 V-Y). This latter expression is possibly associated with the ventral-most tissue of the opisthosomal limb buds.

Compared to *Lbx-A*, expression of *P. tepidariorum Lbx-B* appears later and is restricted to the prosomal appendages (Figure 9 b-d). *Lbx-B* expression within the developing appendages is similar, but not identical, to that of *Lbx-A*; *Lbx-B* is expressed in fewer ventral cells than *Lbx-A*. (Figure 9 e-f). Expression of the single *P. opilio Lbx* gene is similar to that of *P. tepidariorum Lbx-A*, appearing early during development in the mesoderm of the prosomal appendages (Figure 9 g-h), and later extending into ventral ectodermal cells of the limbs (Figure 9 j-k). Unlike the two *P. tepidariorum Lbx* ohnologues, *P. opilio Lbx* is also expressed at the posterior of the developing embryo (asterisks in Figure 9 h-i).

Expression of both ohnologues of *P. tepidariorum NK7* were only detected in relatively late stages of embryogenesis. *NK7-A* is exclusively expressed in the stomodaeum and in a few patches in the head lobes (Figure 9 l-m). *NK7-B* is expressed in three rings in the developing walking legs (Figure 9 n-p). The expression of *P. opilio NK7* combines the expression of both *P. tepidariorum NK7* genes as it is expressed in the head lobes and in three ventral domains in the legs (Figure 9 q-s), again suggesting the subfunctionalisation of the spider *NK7* genes.

## DISCUSSION

### Evolution of the repertoire and organisation of spider homeobox genes before and after the WGD

We detected widespread conserved patterns of homeobox gene content and clusters across spider genomes that were impacted both by tandem duplications and the WGD. Leite et al. (2018) previously discovered that up to 59% of spider homeobox families were duplicated in arachnopulmonates and that most of these were dispersed duplicates rather than closely linked tandem duplicates. However, the few fragmented spider genomes available at the time meant that whether these were ohnologues or tandem duplicates could not reliably be determined. Here we showed that most duplicated homeobox genes are likely ohnologues retained after WGD, and the majority have been retained in all spiders surveyed (Figure 2 A, Table 2). The arachnopulmonate WGD has therefore played a prominent role in shaping the homeobox gene repertoires of spiders.

Duplicated homeobox gene clusters seem to have evolved asymmetrically following the WGD. Between the two ohnologous Hox clusters as well as the two NK, *Irx*, and SINE clusters, one cluster is more conserved in gene content, orientation, and order than the other. This asymmetry could be a result of mechanisms of regulatory subfunctionalisation (e.g., complementary degenerating mutations to regulatory elements (Force et al. 1999)) acting not only at the level of individual genes, but also on homeobox gene clusters.

Contextualising the organisation of the conserved duplicated homeobox gene clusters across entire spider genomes, we found that synteny is largely conserved among the entelegynes (Figure 6). We also observed a few rearrangements of homeobox clusters associated with chromosomal rearrangements (Figure S4 B and C). In *H. graminicola*, Hox cluster B and NK cluster B are both on chromosome 12, one of the two sex chromosomes, while in the other entelegynes, the synteny blocks that make up the *H. graminicola* sex chromosomes are found on two distinct chromosomes, one with Hox cluster B and the other with NK cluster B. This could have arisen by a rearrangement among the sex chromosomes in *H. graminicola* or independent rearrangements in theridiids and araneids. Given the X_1_X_2_ male and X_1_X_1_X_2_X_2_ female sex determination in most entelegynes including the species shown here (Maddison and Leduc-Robert 2013), differential rates of purifying selection on sex chromosomes versus autosomes may impact the extent to which the synteny of these chromosomes is conserved, thus affecting the homeobox clusters on them (Kořínková and Král 2013; Charlesworth 2017; Cordellier et al. 2020).

There seems to be a disconnect between the relative organisation of spider Hox clusters and their regulation. Hox genes exhibit spatial and temporal collinearity (Ferrier 2019 and the references therein). The regulatory elements responsible for temporal collinearity constrain Hox gene organisation, thus it has been hypothesised that Hox cluster integrity is essential for the mechanism of temporal collinearity (Duboule 2007; Krumlauf 2018; Ferrier 2019). While spatial collinearity is exhibited by both spider Hox clusters, temporal collinearity is more evident in cluster A, though this cluster is less organised (Schwager et al. 2017; Turetzek et al. 2022). Hox cluster organisation and regulation vary between animals, for example, in mammals, each Hox cluster is condensed and is regulated globally, while in some invertebrates the Hox cluster is organised and expressed in sub-clusters (Kmita and Duboule 2003; Deschamps and Duboule 2017; Wang et al. 2017). This necessitates a better understanding of the relationship between the organisation and regulation of the ohnologous spider Hox clusters.

The conservation of other homeobox clusters indicates that they may similarly be constrained by regulatory elements. ANTP-class Super-Hox genes arose with the Hox cluster, and many are still linked to one another and to the Hox cluster across bilaterians (Butts et al. 2008; Ferrier 2016b). However, in spiders, the only consistent pattern among Super-Hox genes was the conserved clustering of *Abox* and *Meox*, though these two genes were not linked to either Hox cluster. This suggests that amid many rearrangements and gene losses, there may be a functional constraint on this two-gene cluster in spiders that would not have been recognised without comparisons across this taxonomic breadth.

Our comparison of NK cluster genes in spiders with other bilaterians revealed a pattern of conserved sub-clusters. In spiders, we found a core NK cluster consisting of a subset of the bilaterian NK complement largely conserved in gene order (Figure 4). Its ohnologous cluster, NK cluster B, is less conserved but is linked to more of the bilaterian NK complement (Figure 4). Some NK gene organisation patterns seen in spiders are similar to those in other arthropods. While the NK genes in different drosophilid species have undergone a series of rearrangements, fragmenting and then reuniting the NK cluster, certain gene pairs or triplets are consistently clustered and move as units referred to as “contiguities” (Chan et al. 2015). In spiders, *NK3* and *NK4* are adjacent, as in *D. melanogaster*, *T. castaneum*, *Bombyx terrestris*, many butterflies, and *Anopheles gambiae*, and *Tlx* and *Lbx* are adjacent, as in *D. melanogaster*, *T. castaneum*, several butterflies, *A. gambiae* and *I. scapularis* (Figure 4) (Luke et al. 2003; Chan et al. 2015; Mulhair et al. 2022). Though *NK1* has undergone a recent tandem duplication in spiders, *NK5* and *NK1* are also adjacent in *T. castaneum*, *B. terrestris*, *A. gambiae*, and *I. scapularis*, but not *D. melanogaster* or butterflies (Figure 4) (Chan et al. 2015; Mulhair et al. 2022).

These NK sub-clusters are even conserved across other bilaterians (Luke et al. 2003; Larroux et al. 2007; Wotton et al. 2009; Hui et al. 2012; Mulhair et al. 2022). In the annelid *Platynereis dumerilii* and the invertebrate chordate amphioxus *Branchiostoma floridae*, *NK3* and *NK4* are clustered, as are *Lbx* and *Tlx* (Hui et al. 2012), while in human, one *NK4* ohnologue clusters with *Tlx* and *Lbx* ohnologues (Luke et al. 2003; Wotton et al. 2009). In contrast, the NK sub-clusters of tardigrades consist of different subgroups (Treffkorn et al. 2018), which most likely is a result of rearrangements in this lineage. Overall, these patterns suggest that while the entire NK cluster has undergone many rearrangements in different animal lineages, these events have repeatedly occurred between these sub-clusters, and far less often within them. This semi-flexible organisation distinguishes the NK cluster from the Hox cluster, which has a more consistent gene complement and order, but each of the NK sub-clusters may be maintained by a conserved regulatory mechanism similar to the Hox cluster.

### NK cluster genes show conserved and subfunctionalised expression

We observed that the genes in NK sub-clusters are expressed in similar patterns. NK cluster genes are broadly involved in patterning the mesoderm (Jagla et al. 2001; Saudemont et al. 2008; Treffkorn et al. 2018). The expression of *NK3* and *NK4* in both arachnids is similar to the *D. melanogaster* orthologues *tin* and *bap* in the heart and visceral mesoderm (Azpiazu and Frasch 1993). The fly *Tlx* orthologue, *C15*, and *Lbx* paralogues *lbe* and *lbl* are also expressed in legs like their orthologues in both *P. tepidariorum* and *P. opilio* (Jagla et al. 2001; Campbell 2005; Maqbool et al. 2006). This suggests these genes share conserved cis-regulatory elements that may not only preserve their clustering but also result in their co-expression. It would be interesting to identify these regulatory elements and to investigate the extent of co-expression within NK sub-clusters more broadly across bilaterians.

Though a few examples of potential neofunctionalisation have been described, subfunctionalisation seems to be the more common fate for duplicated homeobox genes (Schwager et al. 2017; Leite et al. 2018), and other genes (Schomburg et al. 2015; Janssen et al. 2021) in spiders. Using a more thorough annotation of NK genes from the harvestman, we were able to determine the relative functional consequences of tandem duplication versus the WGD, which was particularly important for families with ancient tandem duplication events like *Emx* and *Msx*. Expression of *Emx* tandem paralogues were similar while A and B ohnologues have undergone temporal subfunctionalisation (Leite et al. 2018), despite the *Emx* tandem duplication now estimated to predate the WGD. Furthermore, the two *Emx* paralogues in *P. opilio* are expressed in near-identical patterns (Figure 8) (Leite et al. 2018), which suggests that while the tandem duplication gave rise to two conserved paralogues, *Emx* genes did not subfunctionalise until after the WGD. For *Msx*, we found that *P. opilio Msx2* exhibits a similar expression pattern to *Msx2-B*, which now seems more likely to have undergone subfunctionalisation instead of neofunctionalisation. This stresses the importance of understanding the full evolutionary history of a gene family before inferring sub- and/or neofunctionalisation. Furthermore, the roles of the ancestral single copies of *Msx* and *Emx* are not yet known but could be determined with comparisons to earlier-branching lineages of chelicerates or arachnids.

It is thought that duplicated genes encoding developmental transcription factors like homeobox genes are more likely to undergo regulatory subfunctionalisation, driven by the complementary loss of enhancers (Force et al. 1999; Jiménez-Delgado et al. 2009; Espinosa-Cantú et al. 2015; Marlétaz et al. 2018). Indeed, the clear patterns of subfunctionalisation observed for the *P. tepidariorum* NK ohnologues and single-copy orthologues from *P. opilio* despite no NK cluster ohnologues being found to be under positive selection suggests changes primarily occurred in their regulatory regions.

### Comparisons to the outcomes of other WGD events

Our results suggest that there have been similar outcomes between the arachnopulmonate WGD and the likely independent WGDs in horseshoe crabs (Nossa et al. 2014; Kenny et al. 2016; Shingate et al. 2020; Nong et al. 2021). Here, many homeobox gene families are expanded to four or more copies and there are multiple duplicated homeobox clusters, including up to six Hox clusters, six NK clusters, and seven SINE clusters as a result of up to three WGDs (Nong et al. 2021). Like spiders, in horseshoe crabs some ohnologous clusters are more intact than others; for instance, two of the six Hox clusters are intact and ordered, while the others have lost between two and eight genes (Shingate et al. 2020). However, in the absence of additional chromosome-level assemblies, the extent to which this applies to other horseshoe crabs is not yet clear.

In comparison to vertebrate homeobox gene clusters resulting from the 2R WGD, the ohnologous clusters of spiders are more conserved in terms of gene content. Across the four human Hox clusters, complementary gene losses resulted in no cluster containing all gene families, but in spiders, one Hox cluster is intact while the other has lost only two genes (Figure 3). The four ohnologous human NK clusters are spread across five chromosomes, and most genes are retained in two or maximum three copies (Luke et al. 2003; Wotton et al. 2009). While in spiders the HRO cluster is intact, *Hbn* is not found in chordates, and *Otp* is not linked to *Rax* or *Rax2* in the human genome (Mazza et al. 2010). However, for SINE genes, the clusters have evolved similarly between vertebrates and spiders following WGD (Kawakami et al. 2000; Ferrier 2016b). In both of these lineages, linkage of all three SINE genes is conserved on one chromosome, while the ohnologous cluster(s) have undergone losses and rearrangements. Therefore, the pattern of one copy being retained intact while the other has degenerated, as seen for most ohnologous spider homeobox gene clusters, is similar for only some human homeobox clusters. Still, these few similarities illustrate how duplication can lead to asymmetric degradation of otherwise conserved gene clusters.

Besides conserved clusters, some homeobox genes have returned to the single copy in both humans and spiders or scorpions, namely *Hlx*, *Hhex*, *Mnx*, *Dmbx*, *Otp*, *Prop*, and *Mkx* (Holland et al. 2007; Leite et al. 2018). Genes often found in the single copy in both spiders and horseshoe crabs are *Bari*, *Barx*, *CG11294*, *Dmbx*, *Hhex*, *Meox*, *Mkx*, *Msxlx*, and *NK4* (Nong et al. 2021). This suggests that certain genes may be constitutive singletons, disadvantageous in multiple copies, or unable to subfunctionalise and thus lost. However, these comparisons are only between three distantly related species; wider sampling of WGD lineages may reveal clearer patterns of genes that tend to return to the single copy and the underlying reasons.

WGDs are hypothesised to have had important evolutionary consequences, primarily based on the finding that 2R WGD occurred along the vertebrate stem. However, the impact of WGDs in other animal lineages remains unclear. We still do not know to what extent the evolution of the morphological novelties and species diversity of spiders was driven by the arachnopulmonate WGD, though our work contributes to a growing body of evidence detailing widespread subfunctionalisation of developmental ohnologues.

The homeobox genes we have characterised here as well as other developmental genes provide candidates for future investigations, which, in concert with future studies of other chelicerate lineages, advance our aim to understand the impact of the arachnopulmonate WGD and WGDs more widely across animals.

## CONCLUSIONS

Analyses of WGDs in arachnopulmonates, horseshoe crabs, and vertebrates shows that homeobox genes, as well as other regulatory genes are commonly retained in duplicate (Cañestro et al. 2013; Berthelot et al. 2014; Leite et al. 2016; Harper et al. 2021; Janssen et al. 2021). Across eight spiders, representing three major clades we found that the majority of duplicated homeobox genes likely originated in the arachnopulmonate WGD and that often, cluster or sub-cluster integrity has been maintained in at least one ohnologous copy. Comparisons of expression patterns of selected homeobox genes support extensive subfunctionalisation among spider ohnologues. Taken together, our results help us understand the impact of WGD and tandem duplication on the evolution of homeobox genes and genomes in spiders and provide an important comparison to WGD in other animals.

## MATERIALS AND METHODS

### Homeobox gene annotation

We classified the homeobox gene complement of the publicly available chromosome-level genomes from eight spider species, *D. silvatica* (Sánchez-Herrero et al. 2019; Escuer et al. 2022), *D. plantarius* (Blaxter et al. 2022), *L. elegans* (Wang et al. 2022), *P. tepidariorum* (Zhu et al. 2023), *H. graminicola* (Zhu et al. 2022), *A. bruennichi* (Sheffer et al. 2021), *T. antipodiana* (Fan et al. 2021), and *T. clavata* (Hu et al. 2022). We examined the previous homeobox gene classification of the scorpion *C. sculpturatus*, which shared the ancestral WGD, as an outgroup to the spiders (Schwager et al. 2017; Leite et al. 2018). As non-WGD outgroups, we included two arachnids, the harvestman *P. opilio* (Gainett et al. 2021), and the tick *I. scapularis* (De et al. 2023), as well as three mandibulates: the centipede *S. maritima* (Chipman et al. 2014; So et al. 2022), the fruit fly *D. melanogaster* (BDGP6.32), in which many of these homeobox genes and families were first described (Zhong et al. 2008), and the beetle *T. castaneum* (Herndon et al. 2020), which was used by Butts et al. (2008) to describe the Super-Hox cluster. Only chromosome-level assemblies were used for synteny analyses, but homeobox genes were counted and classified in all genomes. Genome accessions are shown in Table 4.

**Table 4.** – Genome assemblies used in this study and accessions. NCBI accession numbers or sequence database DOIs are given for genomes with homeobox genes annotated in this study for the first time. Citations are given for genomes with published homeobox gene annotations. Genome size, N50, and BUSCO scores are shown for each genome as calculated for the annotated chromosome-level assemblies with the arthropoda_odb10 database using BUSCO v5.4.7 or were taken from NCBI and the genome papers (cited in the text above). Abbreviations: C: completeness; S: single copy; D: duplicated. The first-listed *A. bruennichi* genome, sequenced by the Darwin Tree of Life project, had a higher N50 and larger size, so was used for homeobox annotation, however it has not been annotated, so an older version, the second-listed, was used for the whole genome synteny comparison as it has a corresponding annotation. A homeobox annotation was also conducted on this older version and can be found at the bottom of Table S1. The older *P. tepidariorum* assembly was not to chromosome-level so homeobox genes were reannotated on the newer chromosome-level genome despite existing annotations on the previous version.

| Species | Accession or Citation | Size | N50 | BUSCO |
| --- | --- | --- | --- | --- |
| *T. clavata* | https://spider.bioinfotoolkits.net/base/download/-1 | 2.6 Gb | 202.1 Mb | C:83.8%<br>[S:80.8%, D:3.0%] |
| *T. antipodiana* | http://dx.doi.org/10.5524/100868 | 2.3 Gb | 172.9 Mb | C:96.3%<br>[S:92.5%, D:3.8%] |
| *A. bruennichi* (DTOL) | GCF_947563725.1 | 1.8 Gb | 139.1 Mb | unannotated |
| *A. bruennichi* (OLD) | GCA_015342795.1 | 1.7 Gb | 124.2 Mb | C:87.0%<br>[S:81.5%, D:5.5%] |
| *H. graminicola* | GCA_023701765.1 | 936.1 Mb | 77.1 Mb | C:96.0%<br>[S:92.2%, D:3.8%] |
| *P. tepidariorum* (OLD) | GCF_000365465.2; annotated by Schwager et al. 2017, Leite et al. 2018 | 1.4 Gb | 4.1 Mb | 98% completeness reported |
| *P. tepidariorum* (NEW) | https://doi.org/10.11922/sciencedb.o00019.00014 | 1.1 Gb | 93.9 Mb | C:95.2%<br>[S:92.4%, D:2.8%] |
| *L. elegans* | http://dx.doi.org/10.5524/102210 | 1.6 Gb | 114.3 Mb | C:58.3%<br>[S:55.2%, D:3.1%] |
| *D. plantarius* | GCA_907164885.2 | 2.8 Gb | 216.7 Mb | unannotated |
| *D. silvatica* | GCA_006491805.2 | 1.4 Gb | 174.2 Mb | C:77.2%<br>[S:74.7%, D:2.5%] |
| *C. sculpturatus* | GCA_000671375.2; annotated by Schwager et al. 2017, Leite et al. 2018 | 925.5 Mb | 537.5 kb | 96.8% completeness reported |
| *P. opilio* | GCA_019434445.1 | 576.9 Mb | 211 kb | 95.1% reported |
| *I. scapularis* | GCA_016920785.2; annotated by Schwager et al. 2017, Leite et al. 2018 | 2.2 Gb | 132.1 Mb | C:98.8%<br>[S:93.4%, D:5.4%] |
| *S. maritima* | GCA_000239455.1; annotated by Chipman et al. 2014, Leite et al. 2018 | 176.2 Mb | 139.5 kb | 93.8% completeness reported |
| *T. castaneum* | GCA_000002335.3; HomeoDB; Zhong et al. 2008, Zhong and Holland, 2011 | 165.9 Mb | 4.5 Mb | C:98.4%<br>[S:97.7%, D:0.7%] |
| *D. melanogaster* | GCA_000001215.4; HomeoDB; Zhong et al. 2008, Zhong and Holland, 2011 | 143.7 Mb | 25.3 Mb | C:92.9%<br>[S:92.3%, D:0.6%] |

We used tBLASTn searches (Altschul et al. 1990) with the contents of HomeoDB (accessed July 2020) (Zhong et al. 2008; Zhong and Holland 2011), as well as the previous classification of *P. tepidariorum* homeobox genes (Schwager et al. 2017; Leite et al. 2018) as the query set to retrieve homeodomain sequences from all eight spider genomes, and re-annotated the homeobox genes for the harvestman, tick, and centipede. Homeodomain sequences were then curated by tBLASTn searches against annotated gene models or transcriptomes for all species except *D. plantarius*, for which an annotation is not available, and classified based on sequence similarity and diagnostic residues of the homeodomain for all families. For some gene families with several duplication events, *e.g.*, *Irx*, *Pax4/6*, *Pax3/7*, as well as for the NK cluster genes, phylogenies were constructed (see below). All homeodomain sequences can be found in the supplementary information (Table S1).

### Phylogenetics

To determine the timing of duplication of the arachnopulmonate *Irx* genes relative to pre-duplicate paralogues, we made a full-length amino acid sequence alignment from a from a subset of our spider species, the scorpion, and a transcriptome from another synspermiatan spider, *Segestria senoculata* (Baudouin-Gonzalez unpublished). Full sequences for genomes with poor annotations were curated from GenBank searches and BLASTp (Altschul et al. 1990) searches against nr (Benson et al. 2018). These sequences were aligned with MUSCLE set to the protein alignment defaults (Edgar 2004) and manually curated and trimmed to remove gaps. We inferred a Maximum-Likelihood phylogeny with IQ-TREE (Nguyen et al. 2015), determined the best substitution model with ModelFinder (Kalyaanamoorthy et al. 2017), and used 1,000 bootstrap replicates to optimise topology support. The phylogeny was visualised with FigTree (FigTree 2018). The full and trimmed alignments and the phylogeny can be found in the supplementary information on GitHub (Figure S1, [https://github.com/madeleineaaseremedios/SpiderHomeoboxSequences]). Other family phylogenies were made using only the homeodomain sequence and aligned manually, but inferred as for the *Irx* family tree, except for the NK cluster and NK-linked genes phylogeny, which used full protein sequences and was aligned with MUSCLE set to the protein default (Edgar 2004) (Figure S2).

### Whole genome synteny comparison

We performed whole genome synteny comparisons to explore chromosomal rearrangements that might underlie trends in homeobox placement in spider genomes. We used all spider species mentioned above, except for *D. plantarius* due to the lack of genome annotation, and compared these to non-spider species *I. scapularis*, *T. castaneum*, and *D. melanogaster*. Hierarchical orthogroups (HOGs) of all non-homeobox genes were identified using Orthofinder (Emms and Kelly 2019) with default settings, specifying the species tree: ((Dmel,Tcas)N1,(Isca,(Dsil,((Ptep,Lele)N5,(Hgra,(Abru,(Tcla,Tant)N8)N7) N6)N4)N3)N2)N0);. These were combined with HOGs of our homeobox orthology and used to reconstruct ancestral gene orders with Agora (Muffato et al. 2023), using the basic pipeline. However, we used Agora to illustrate chromosomes with macrosynteny and orthology relationships, rather than the ancestral synteny blocks, termed Contiguous Ancestral Regions (CARs) detected by Agora, because it resolved a better signal of chromosomal rearrangements across the whole genome compared to the smaller coverage of CARs. We also used MCScanX (Wang et al. 2012) to capture microsynteny relationships between spider genomes and used Circos (Krzywinski et al. 2009) to plot synteny relationships and homeobox positions in the genome.

### Coding sequence evolutionary analyses

For core NK cluster genes, we curated full coding sequences for a subset of species with the best annotations: *P. tepidariorum*, *A. bruennichi*, *D. silvatica*, *D. melanogaster*, *T. castaneum* and *I. scapularis*, as well as four species not included in the synteny comparison: the centipede *S. maritima*, the harvestman *P. opilio*, the scorpion *C. sculpturatus*, and the onychophoran *Euperipatoides rowelli*, for which NK homeobox genes have been described, but which lack chromosomal-level genome assemblies (Chipman et al. 2014; Schwager et al. 2017; Leite et al. 2018; Treffkorn et al. 2018). The NK cluster genes contained families with different duplication histories, including WGD ohnologues in arachnopulmonates, tandem duplications both before and after the WGD, and families with only single genes represented after the WGD, allowing us to compare possible signatures of subfunctionalisation between the different evolutionary trajectories and different types of duplicates.

We used aBSREL v2.3 (Smith et al. 2015) to detect selection on each branch of the phylogeny for each of the spider NK core cluster gene families (*Msx*, *NK3*, *NK4*, *Tlx*, *Lbx*, and *NK7*) and *Emx*. We used sequences from three spiders, *P. tepidariorum*, *D. silvatica*, and *A. bruennichi*, and the scorpion, harvestman, tick, fly, and beetle, as above, and rooted the trees with sequences from the onychophoran (Treffkorn et al. 2018). We created and trimmed codon alignments with PAL2NAL (Suyama et al. 2006) from coding sequences and protein sequence alignments inferred with MUSCLE set to the protein alignment default (Edgar 2004). We ran aBSREL using hyphy v2.5.46 and the outputs were visualised with HyPhy Vision at [http://vision.hyphy.org/] (Pond et al. 2005; Kosakovsky Pond et al. 2020). We also conducted pairwise comparisons of dN/dS ratios between *P. opilio* and *P. tepidariorum* for NK cluster genes calculated with codeML (Goldman and Yang 1994) from codon alignments inferred as above for each gene pair. Sequence data and aBSREL outputs can be found in the supplementary information on GitHub at [https://github.com/madeleineaaseremedios/NKseqevoldata] and visualised with HyPhy Vision as above.

### Animal husbandry, embryo collection, PCR, riboprobe synthesis, and in situ hybridization

Embryos of the spider *P. tepidariorum* were collected from the colony established in Uppsala, Sweden, and maintained as described in Prpic et al. (2008). Embryos of the harvestman *P. opilio* were collected from wild caught specimens in Uppsala, Sweden. Adult specimens and embryos of *P. opilio* were treated as described in Janssen et al. (2021). We applied the staging system of Mittmann and Wolff (2012) for *P. tepidariorum*, and Gainett et al. (2022) for *P. opilio*. We investigated all developmental stages of *P. tepidariorum* and *P. opilio* from the formation of the early germ band to dorsal closure. Total RNA from *P. tepidariorum* and *P. opilio* was isolated from a mix of embryonic stages using TRIzol (Invitrogen). mRNA was isolated from total RNA using the Dynabeads mRNA Purification System (Invitrogen) and reverse transcribed into cDNA using the SuperScriptII system (Invitrogen). PCRs were performed applying sets of gene specific primers. For all investigated genes, reverse primers were ordered with additional 5-prime T7-promotor sequences (gggTAATACGACTCACTATAG) for subsequent antisense RNA probe synthesis (David and Wedlich 2001). *In situ* hybridizations were performed as per Janssen et al. (2018). Primer sequences can be found in Table S2.

## DATA AVAILABILITY STATEMENT

All data underpinning this study is available through the links above or in the supplementary information.

## AUTHOR CONTRIBUTIONS

APM and LSR conceived of the project. MEAR annotated the homeobox genes, conducted the phylogenetics and protein sequence analyses, and drafted the manuscript. RJ characterised the developmental expression of NK genes. DJL conducted the whole genome macrosynteny analysis. All authors contributed to writing and approved the final version of the manuscript.

## ACKNOWLEDGEMENTS

We would like to thank Julio Rozas and Paula Escuer for early access to the *D. silvatica* genome. This project was funded in by a NERC grant (NE/T006854/1) to APM and LSR.

