## Supplementary material for "Evolution of the spider homeobox gene repertoire by tandem and whole genome duplication": Figures S1-5 and Tables S1-3: FigureS1-5andTableS1-2text.pdf

**Supplementary Figure 1 – Maximum-likelihood phylogeny of arthropod *Irx* genes.** Branch lengths are substitutions per site. Branch support values are % of 1,000 ultrafast bootstrap replicates. The tree is rooted arbitrarily between the *mirr/Irx2* clade and the *Irx3* and *ara/caup/Irx1* clades. Species abbreviations are: Abru *Argiope bruennichi*, Ptep *Parasteatoda tepidariorum*, Cscu *Centruroides sculpturatus*, Mmar *Mesobuthus martensii*, Dsil *Dysdera silvatica*, Ssen *Segestria senoculata*, Popi *Phalangium opilio*, Isca *Ixodes scapularis*, Tcor *Trigoniulus corallinus*, Smar *Strigamia maritima*, Dmel *Drosophila melanogaster*, Tcas *Tribolium castaneum*. Sequence IDs and accessions are shown after the species name; manual means curated manually with tBLASTn searches against the genome or transcriptome to annotate a fuller sequence.

**Supplementary Figure 2 – Maximum-likelihood phylogeny of panarthropod NK cluster genes.** Branch lengths are substitutions per site. Branch support values are % of 1,000 ultrafast bootstrap replicates. The tree is rooted with *D. melanogaster lab*. Species abbreviations are: Dmel *Drosophila melanogaster*, Tcas *Tribolium castaneum*, Smar *Strigamia maritima*, Erow *Euperipatoides rowelli*, Isca *Ixodes scapularis*, Cscu *Centruroides sculpturatus*, Ptep *Parasteatoda tepidariorum*, Abru *Argiope bruennichi*, Dsil *Dysdera silvatica*, Popi *Phalangium opilio*. Sequence IDs and accessions are shown after the species name; manual means curated manually with tBLASTn searches against the genome or transcriptome to annotate a fuller sequence.

The alignments can be accessed at:
[https://github.com/madeleineaaseremedios/SpiderHomeoboxSequences].

**Supplementary Figure 3 – Distribution of all homeodomain sequences across eight spider and four outgroup genomes.** A) *D. melanogaster* B) *T. castaneum* C) *S. maritima* D) *I. scapularis* E) *D. silvatica* F) *D. plantarius* G) *L. elegans* H) *P. tepidariorum* I) *H. graminicola* J) *A. bruennichi* (v2) K) *T. antipodiana* L) *T. clavata*. Chromosomes are grey bars, drawn to length scale. Genes annotated to the left are on the reverse strand, and genes on the right are on the forward strand, with 0 Mb at the bottom. Chromosome number, or for *I. scapularis* and *S. maritima*, scaffold accession number are denoted below.

**Supplementary Figure 4 – Circle plot of conserved macrosynteny among chromosome-level entelegyne genomes.** A) A circular version of the plot shown in Figure 6 highlights conserved syntenic blocks between the chromosomes of *H. graminicola* and the other entelegyne species. Each *H. graminicola* chromosome is individually coloured and positions of homeobox clusters are labeled. Each chromosome in the other species is painted respective of its synteny relationships to *H. graminicola*, and the positions of all identified homeobox genes are shown as black lines outside of the chromosomes. B) A focused plot showing only the X1X2 chromosomes reveals the rearrangements associated with the dispersal of NK-B and HOX-B relative to *H.*

*graminicola*. C) A focused plot shows the translocation of part of one chromosome in *T. antipodiana* containing NK cluster A onto another chromosome.

**Supplementary Figure 5 – Expression of *Glomeris marginata NK3*.** In all panels, anterior is to the left and ventral views. The embryo in panel D has been flat-mounted. Developmental staging is indicated in the upper left corner of each panel; the staging system follows Janssen et al. (2004).

Primers:
```
Gm_NK3fw1 GTTGGGTGGCTCGCAACTAA
Gm_NK3fw2 GGAACACCAAAACCTCGCAA
Gm_NK3bw1 gggTAATACGACTCACTATAGGCATGGTTTTGAAACTGTATGCCTT
Gm_NK3bw2 gggTAATACGACTCACTATAGATGGAAATTGCAAGTATTCGACAGA
```

Description:
Early during development, *NK3* is expressed in the form of broad transverse segmental stripes that are strongest in newly formed posterior segments (Figure S3 A, B, D, filled circles). Within the head, strongest expression is in the mandibular segment, and later in the mandibles (Figure S3A, arrowheads). Later during development, expression appears in the head lobes (Figure S3B-D, asterisks), and around the stomodaeum (Figure S3B-D, double-arrowheads). Segmental expression transforms into a salt-and-pepper pattern (Figure S3B) that is reduced to cells of the trunk at stage 4 (Figure 3B-D). This pattern is similar to that of the visceral mesoderm marker *FoxF* in *Glomeris* (Janssen et al. 2022). During limb-bud development, *NK3* is strongly expressed in the mesoderm which is most prominent in the mandibles (Figure S3 C, D, arrows). Note the inlay in D showing a magnification of the left mandible as in panel D. Note absence of *NK3*-positive mesoderm in the antennae and the maxillae.

**Supplementary Table 1 – Homeodomain sequences and genomic positions of homeobox genes in arthropod genomes.** Gene/protein models are provided if they exist. Bilaterian family refers to naming on HomeoDB (Zhong et al. 2008; Zhong and Holland 2011); arthropod families are newly indicated here, based on ancient shared duplications.

**Supplementary Table 2 – Primer sequences for *in situ* hybridisation of NK cluster genes in *P. tepidariorum* and *P. opilio*.**

**Supplementary Table 3 – Substitutions and substitution rates calculated with codeML for pairwise comparisons of NK cluster genes between *P. opilio* and *P. tepidariorum*.**
