## Supplementary material for "Evolution of the spider homeobox gene repertoire by tandem and whole genome duplication": Figures S1-5 and Tables S1-3: FigureS3.pdf

A)

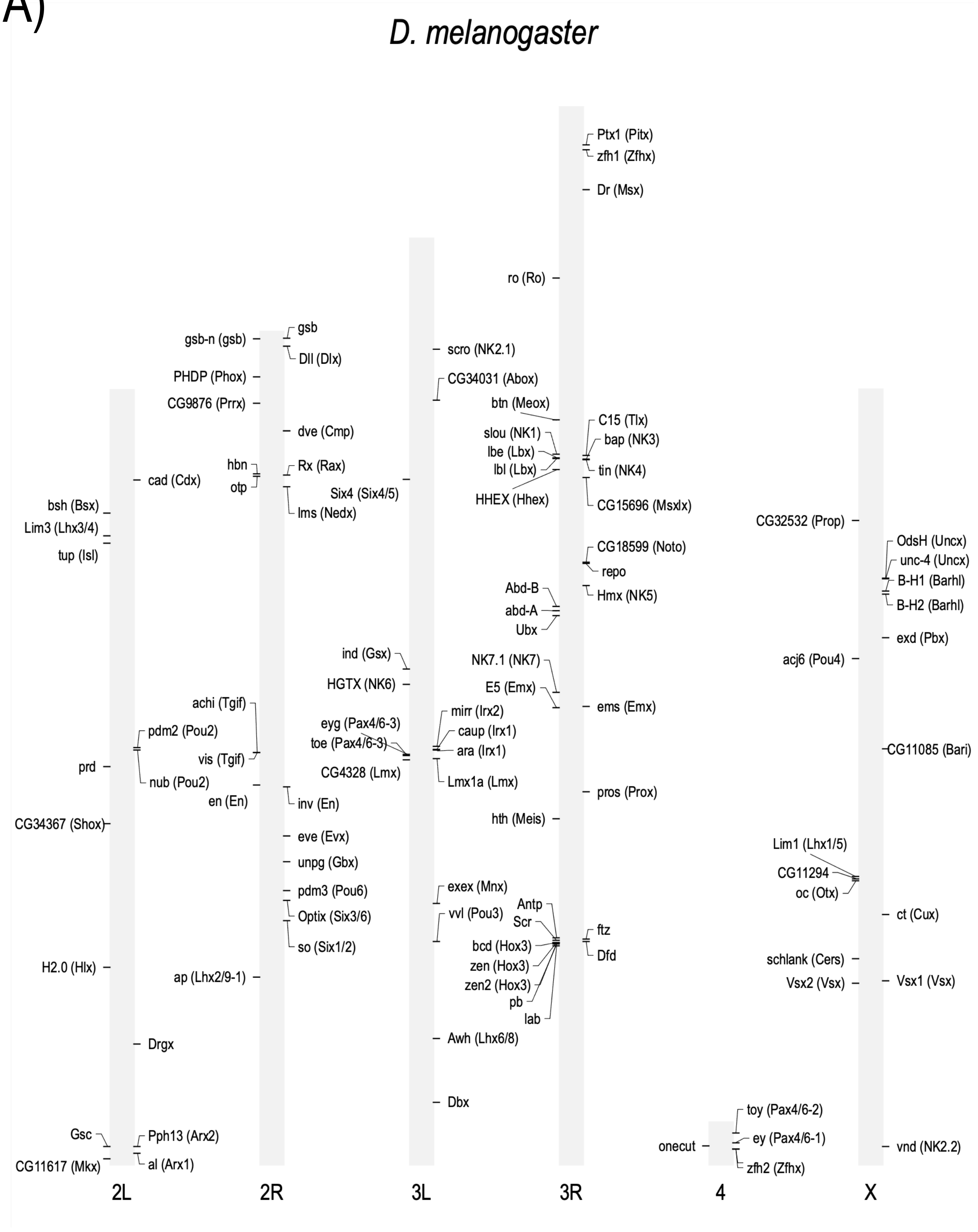
D. melanogaster
Ptx1 (Pitx)
zfh1 (Zfhx)
Dr (Msx)
ro (Ro)
gsb-n (gsb)
gsb
Dll (Dlx)
scro (NK2.1)
PHDP (Phox)
CG34031 (Abox)
CG9876 (Prrx)
btn (Meox)
dve (Cmp)
C15 (Tlx)
slou (NK1)
bap (NK3)
lbe (Lbx)
lbl (Lbx)
tin (NK4)
hbn
Rx (Rax)
otp
cad (Cdx)
Six4 (Six4/5)
HHEX (Hhex)
CG15696 (Msxlx)
bsh (Bsx)
lms (Nedx)
Lim3 (Lhx3/4)
CG32532 (Prop)
tup (Isl)
OdsH (Uncx)
CG18599 (Noto)
unc-4 (Uncx)
repo
B-H1 (Barhl)
Abd-B
Hmx (NK5)
abd-A
B-H2 (Barhl)
Ubx
exd (Pbx)
ind (Gsx)
acj6 (Pou4)
NK7.1 (NK7)
HGTX (NK6)
E5 (Emx)
achi (Tgif)
ems (Emx)
mirr (Irx2)
pdm2 (Pou2)
eyg (Pax4/6-3)
caup (Irx1)
toe (Pax4/6-3)
ara (Irx1)
CG11085 (Bari)
vis (Tgif)
prd
CG4328 (Lmx)
Lmx1a (Lmx)
nub (Pou2)
pros (Prox)
en (En)
inv (En)
hth (Meis)
CG34367 (Shox)
eve (Evx)
Lim1 (Lhx1/5)
unpg (Gbx)
CG11294
oc (Otx)
exex (Mnx)
pdm3 (Pou6)
Antp
ct (Cux)
Optix (Six3/6)
vvl (Pou3)
Scr
ftz
so (Six1/2)
bcd (Hox3)
Dfd
zen (Hox3)
schlank (Cers)
H2.0 (Hlx)
ap (Lhx2/9-1)
zen2 (Hox3)
Vsx2 (Vsx)
Vsx1 (Vsx)
pb
lab
Awh (Lhx6/8)
Drgx
Dbx
toy (Pax4/6-2)
Gsc
Pph13 (Arx2)
onecut
ey (Pax4/6-1)
vnd (NK2.2)
CG11617 (Mkx)
al (Arx1)
zfh2 (Zfhx)
2L
2R
3L
3R
4
X

B)

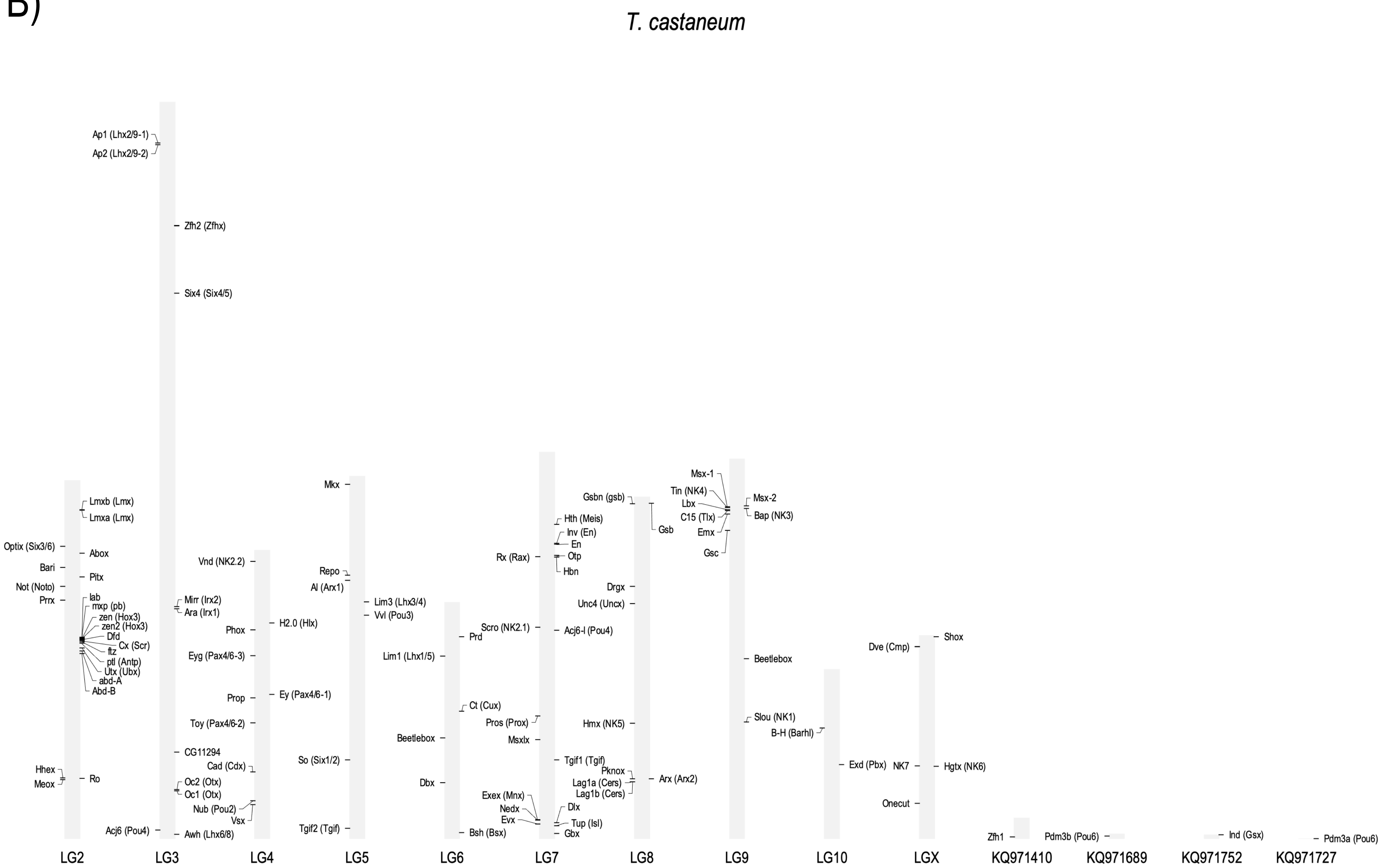
T. castaneum
Ap1 (Lhx2/9-1)
Ap2 (Lhx2/9-2)
Zfh2 (Zfhx)
Six4 (Six4/5)
Lmxb (Lmx)
Lmxa (Lmx)
Optix (Six3/6)
Abox
Bari
Pitx
Not (Noto)
Prrx
lab
mxp (pb)
zen (Hox3)
zen2 (Hox3)
Dfd
Cx (Scr)
ftz
ptl (Antp)
Utx (Ubx)
abd-A
Abd-B
Hhex
Meox
Ro
Acj6 (Pou4)
Vnd (NK2.2)
Mirr (Irx2)
Ara (Irx1)
Phox
Eyg (Pax4/6-3)
Prop
Toy (Pax4/6-2)
CG11294
Cad (Cdx)
Oc2 (Otx)
Oc1 (Otx)
Nub (Pou2)
Vsx
Awh (Lhx6/8)
H2.0 (Hlx)
Ey (Pax4/6-1)
Mkx
Repo
Al (Arx1)
So (Six1/2)
Tgif2 (Tgif)
Lim3 (Lhx3/4)
Vvl (Pou3)
Lim1 (Lhx1/5)
Beetlebox
Dbx
Prd
Ct (Cux)
Bsh (Bsx)
Rx (Rax)
Scro (NK2.1)
Pros (Prox)
Msxlx
Exex (Mnx)
Nedx
Evx
Hth (Meis)
Inv (En)
En
Otp
Hbn
Acj6-I (Pou4)
Tgif1 (Tgif)
Dlx
Tup (Isl)
Gbx
Gsbn (gsb)
Drgx
Unc4 (Uncx)
Hmx (NK5)
Pknox
Lag1a (Cers)
Lag1b (Cers)
Gsb
Arx (Arx2)
Msx-1
Tin (NK4)
Lbx
C15 (Tlx)
Emx
Gsc
Msx-2
Bap (NK3)
Beetlebox
Slou (NK1)
B-H (Barhl)
Exd (Pbx)
Dve (Cmp)
NK7
Onecut
Shox
Hgtx (NK6)
Zfh1
Pdm3b (Pou6)
Ind (Gsx)
Pdm3a (Pou6)
LG2
LG3
LG4
LG5
LG6
LG7
LG8
LG9
LG10
LGX
KQ971410
KQ971689
KQ971752
KQ971727

C)

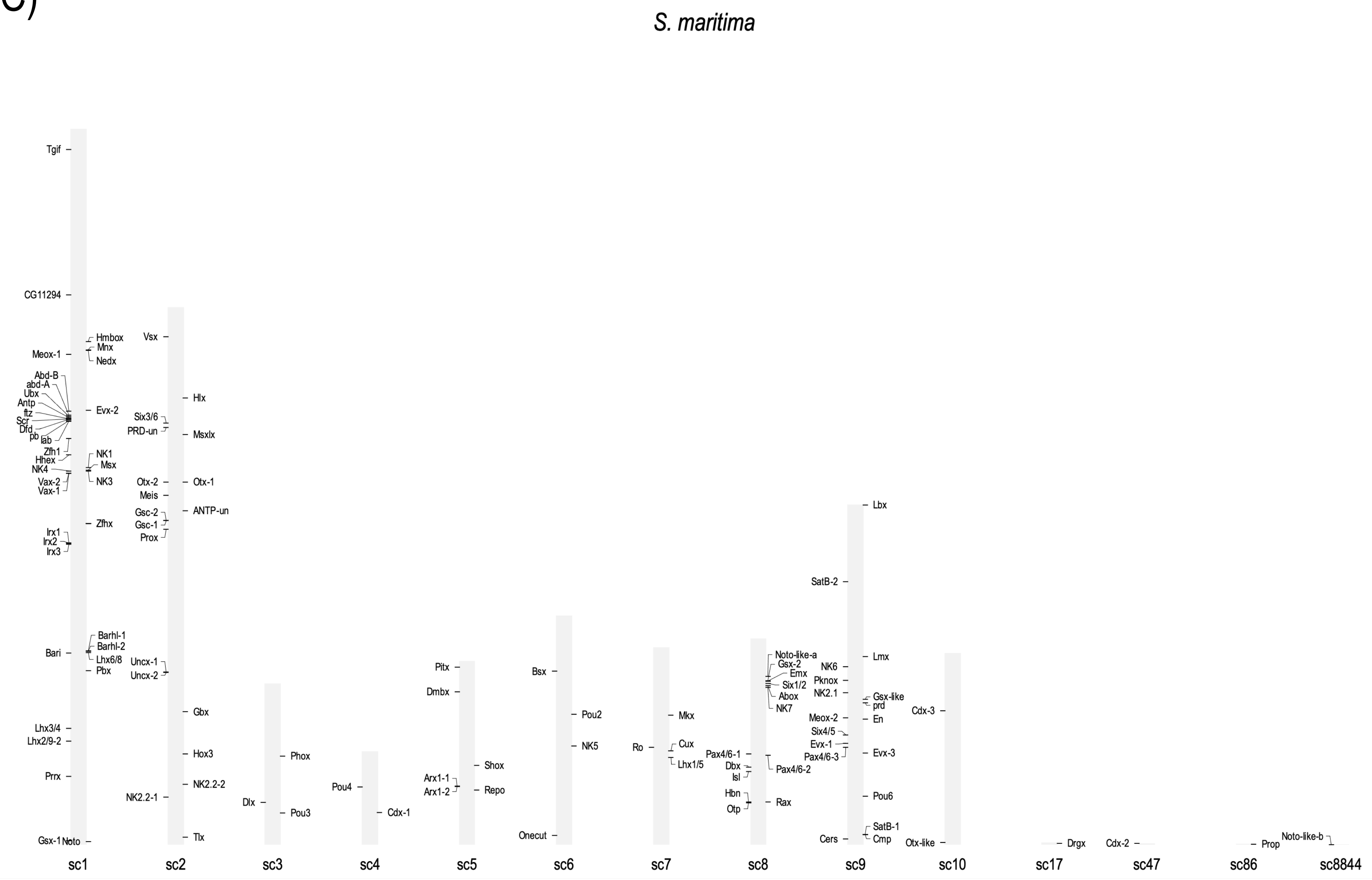
S. maritima
sc1
Tgif
CG11294
Hmbox
Mnx
Meox-1
Nedx
Abd-B
abd-A
Ubx
Antp
ftz
Scr
Dfd
pb
lab
Evx-2
Zfh1
Hhex
NK4
NK1
Msx
NK3
Vax-2
Vax-1
Zfhx
Irx1
Irx2
Irx3
Barhl-1
Barhl-2
Bari
Lhx6/8
Pbx
Lhx3/4
Lhx2/9-2
Prrx
Gsx-1
Noto
sc2
Vsx
Hlx
Six3/6
PRD-un
Msxlx
Otx-2
Otx-1
Meis
Gsc-2
ANTP-un
Gsc-1
Prox
Uncx-1
Uncx-2
Gbx
Hox3
NK2.2-2
NK2.2-1
Tlx
sc3
Phox
Dlx
Pou3
sc4
Pou4
Cdx-1
sc5
Pitx
Dmbx
Shox
Arx1-1
Arx1-2
Repo
sc6
Bsx
Pou2
NK5
Onecut
sc7
Mkx
Ro
Cux
Lhx1/5
sc8
Noto-like-a
Gsx-2
Emx
Six1/2
Abox
NK7
Pax4/6-1
Dbx
Isl
Pax4/6-2
Hbn
Otp
Rax
sc9
Lbx
SatB-2
Lmx
NK6
Pknox
NK2.1
Gsx-like
prd
Meox-2
En
Six4/5
Evx-1
Pax4/6-3
Evx-3
Pou6
SatB-1
Cers
Cmp
sc10
Cdx-3
Otx-like
sc17
Drgx
sc47
Cdx-2
sc86
Prop
sc8844
Noto-like-b

D)

*I. scapularis*

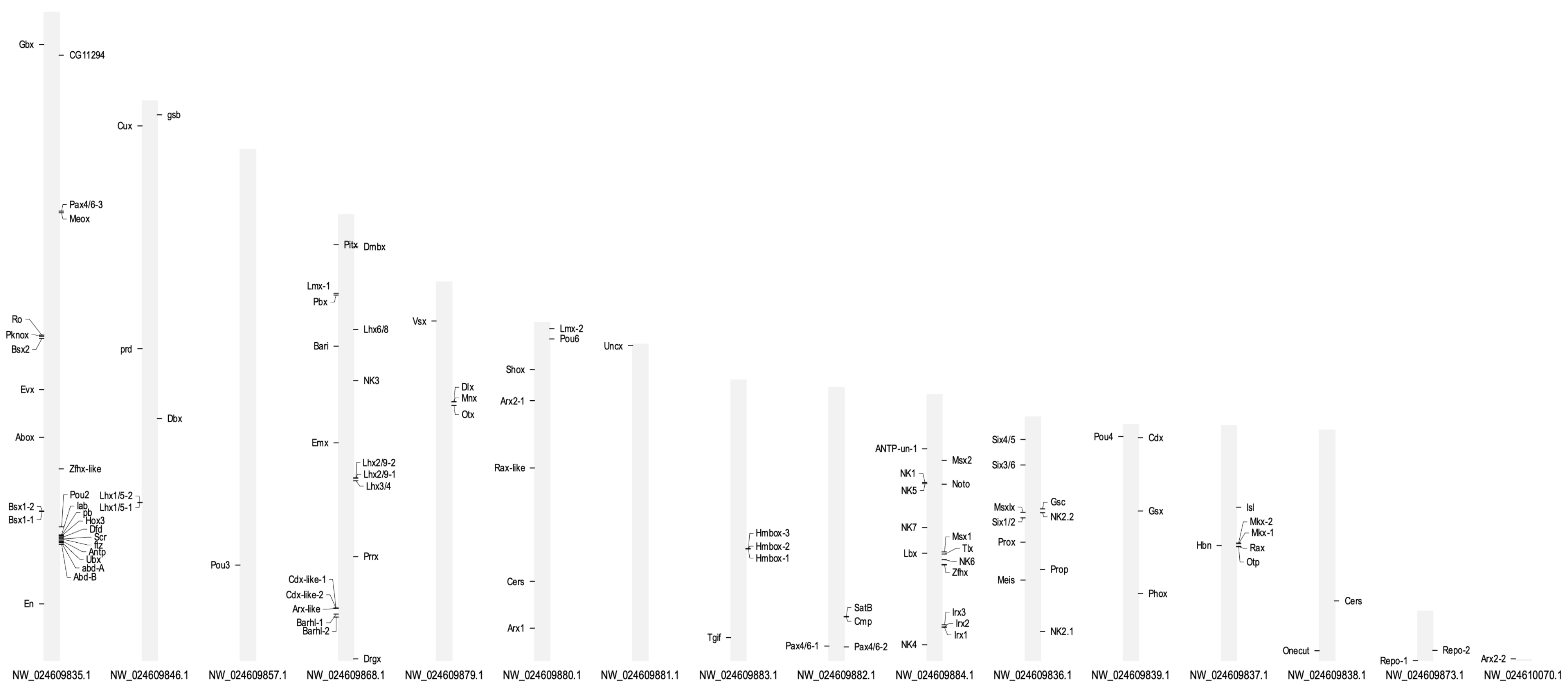

E)

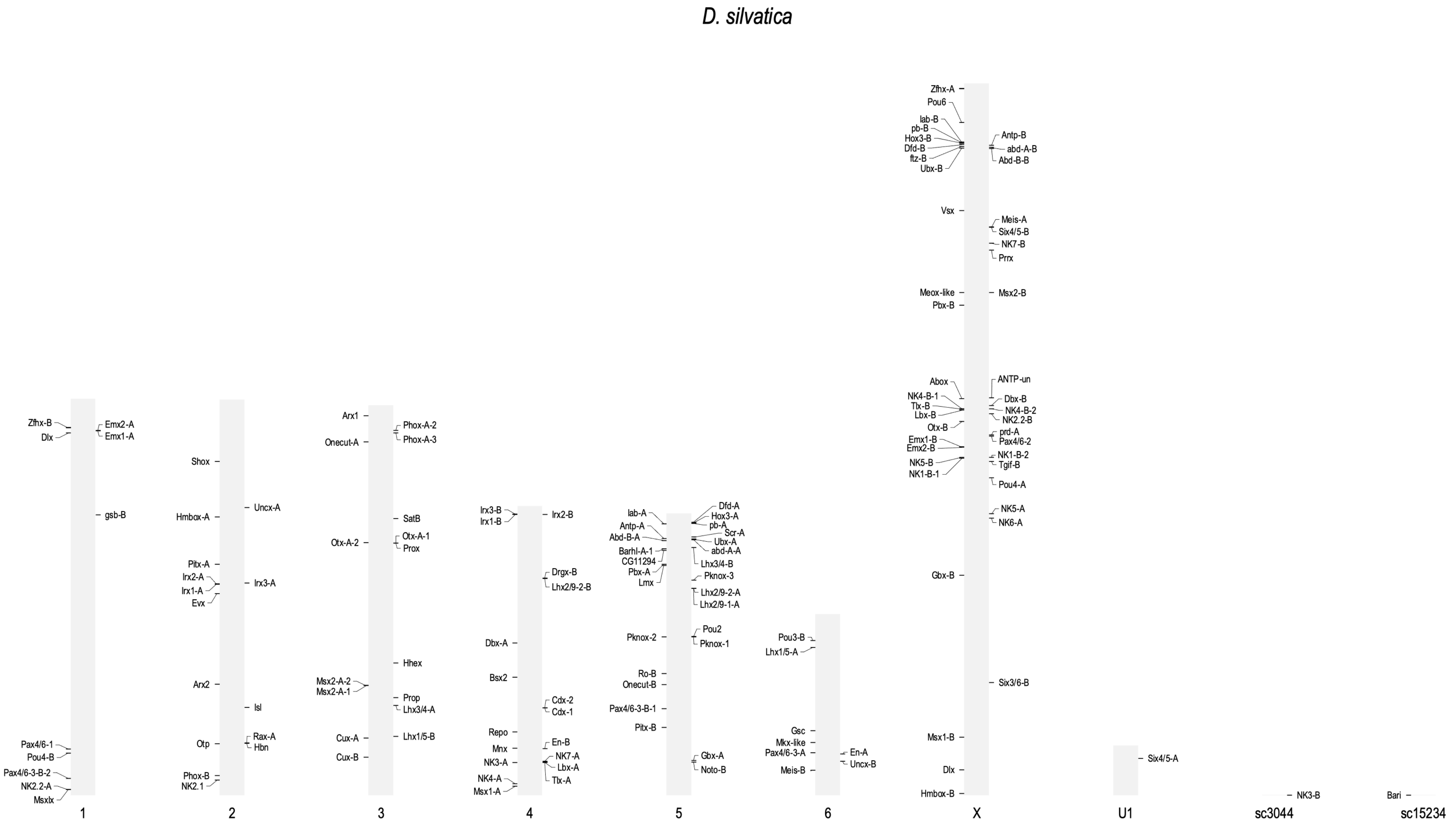
D. silvatica
1
Zfhx-B
Dlx
Emx2-A
Emx1-A
gsb-B
Pax4/6-1
Pou4-B
Pax4/6-3-B-2
NK2.2-A
Msxlx
2
Shox
Hmbox-A
Pitx-A
Irx2-A
Irx1-A
Evx
Arx2
Otp
Phox-B
NK2.1
Uncx-A
Irx3-A
Isl
Rax-A
Hbn
3
Arx1
Onecut-A
Otx-A-2
Msx2-A-2
Msx2-A-1
Cux-A
Cux-B
Phox-A-2
Phox-A-3
SatB
Otx-A-1
Prox
Hhex
Prop
Lhx3/4-A
Lhx1/5-B
4
Irx3-B
Irx1-B
Dbx-A
Bsx2
Repo
Mnx
NK3-A
NK4-A
Msx1-A
Irx2-B
Drgx-B
Lhx2/9-2-B
Cdx-2
Cdx-1
En-B
NK7-A
Lbx-A
Tlx-A
5
lab-A
Antp-A
Abd-B-A
Barhl-A-1
CG11294
Pbx-A
Lmx
Pknox-2
Ro-B
Onecut-B
Pax4/6-3-B-1
Pitx-B
Dfd-A
Hox3-A
pb-A
Scr-A
Ubx-A
abd-A-A
Lhx3/4-B
Pknox-3
Lhx2/9-2-A
Lhx2/9-1-A
Pou2
Pknox-1
Gbx-A
Noto-B
6
Pou3-B
Lhx1/5-A
Gsc
Mkx-like
Pax4/6-3-A
Meis-B
En-A
Uncx-B
X
Zfhx-A
Pou6
lab-B
pb-B
Hox3-B
Dfd-B
ftz-B
Ubx-B
Vsx
Meox-like
Pbx-B
Abox
NK4-B-1
Tlx-B
Lbx-B
Otx-B
Emx1-B
Emx2-B
NK5-B
NK1-B-1
Gbx-B
Msx1-B
Dlx
Hmbox-B
Antp-B
abd-A-B
Abd-B-B
Meis-A
Six4/5-B
NK7-B
Prrx
Msx2-B
ANTP-un
Dbx-B
NK4-B-2
NK2.2-B
prd-A
Pax4/6-2
NK1-B-2
Tgif-B
Pou4-A
NK5-A
NK6-A
Six3/6-B
U1
Six4/5-A
sc3044
NK3-B
sc15234
Bari

F)

*D. plantarius*

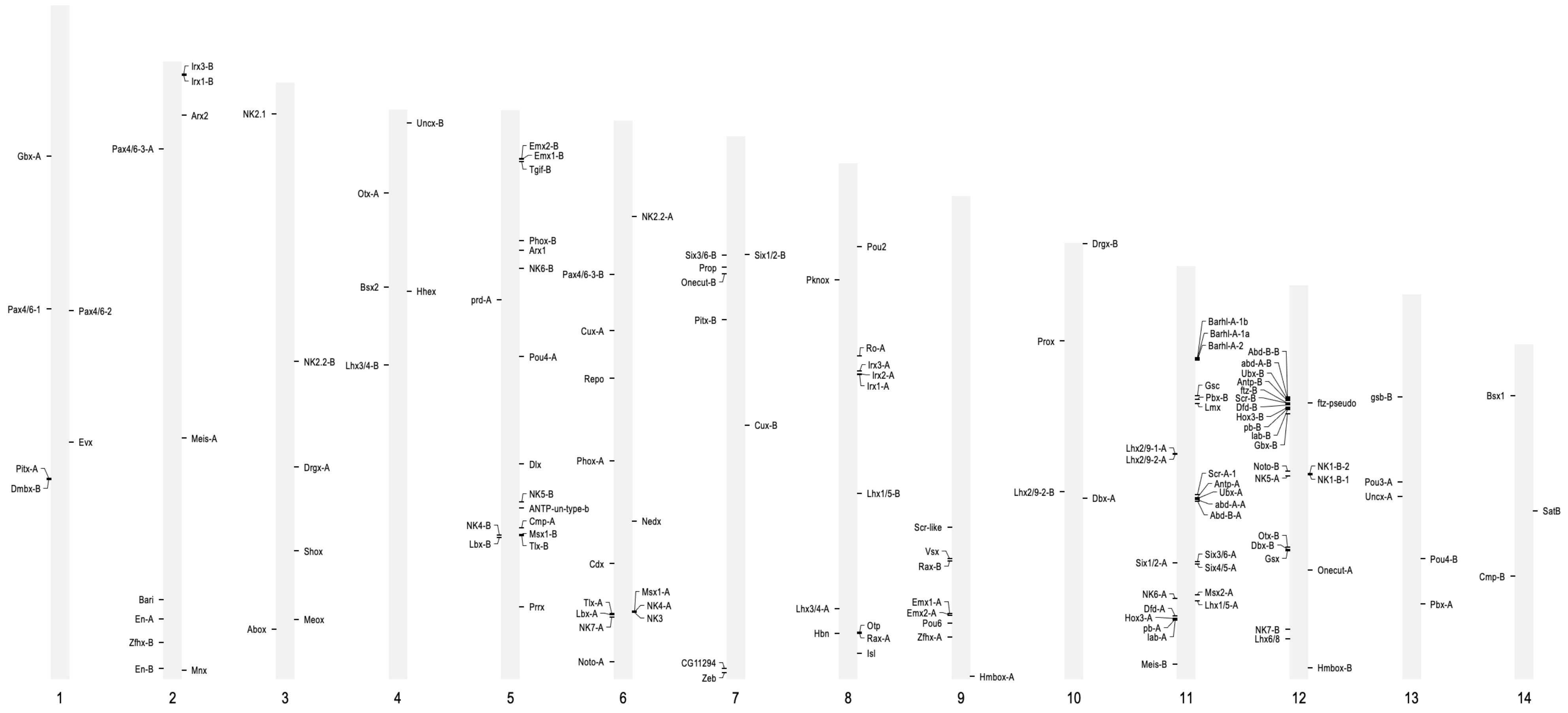

G)

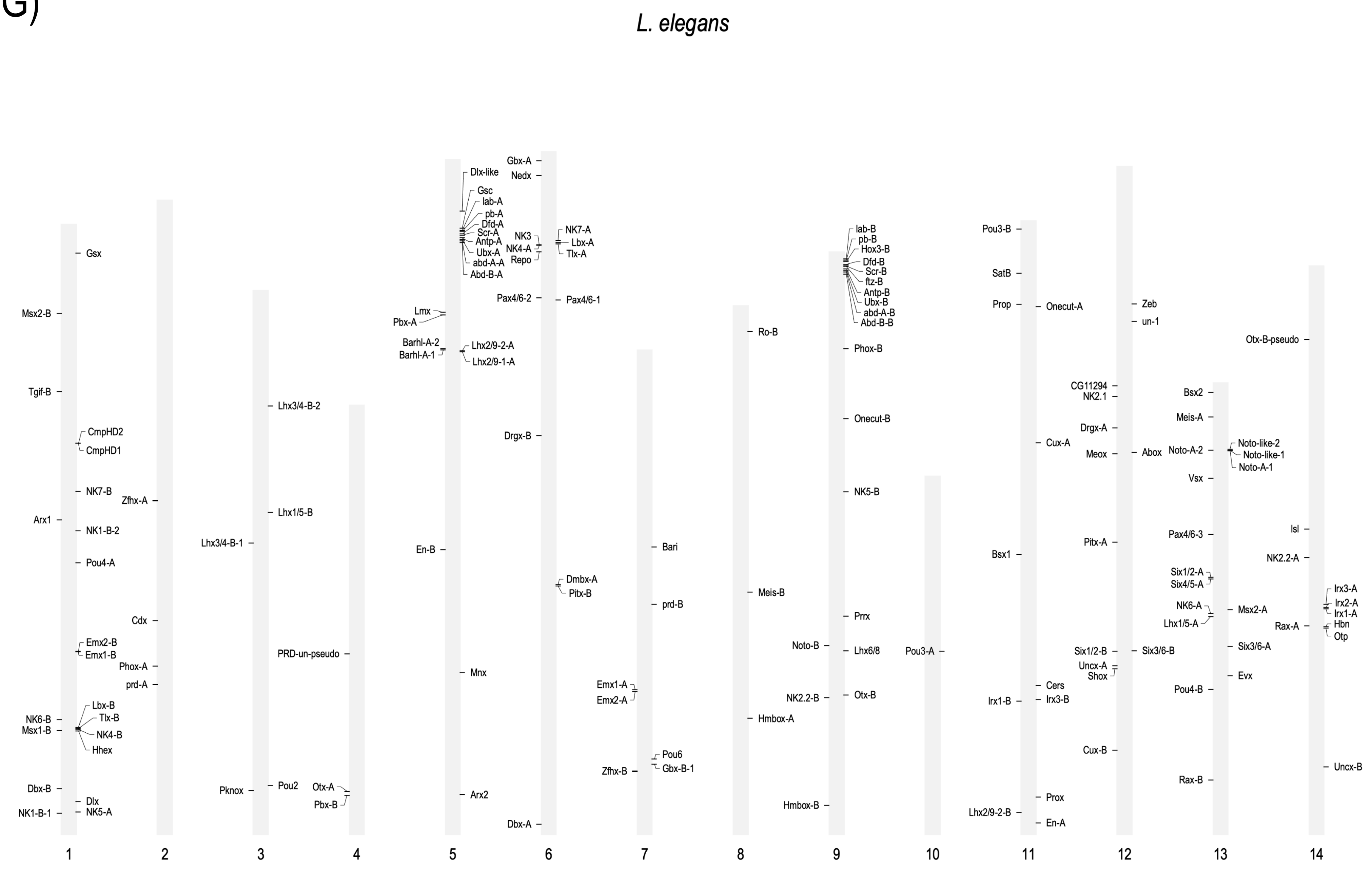
L. elegans
1
Gsx
Msx2-B
Tgif-B
CmpHD2
CmpHD1
NK7-B
Arx1
NK1-B-2
Pou4-A
Emx2-B
Emx1-B
NK6-B
Msx1-B
Lbx-B
Tlx-B
NK4-B
Hhex
Dbx-B
Dlx
NK1-B-1
NK5-A
2
Zfhx-A
Cdx
Phox-A
prd-A
3
Lhx3/4-B-2
Lhx1/5-B
Lhx3/4-B-1
Pknox
Pou2
4
PRD-un-pseudo
Otx-A
Pbx-B
5
Dlx-like
Gsc
lab-A
pb-A
Dfd-A
Scr-A
Antp-A
Ubx-A
abd-A-A
Abd-B-A
Lmx
Pbx-A
Barhl-A-2
Barhl-A-1
Lhx2/9-2-A
Lhx2/9-1-A
En-B
Mnx
Arx2
6
Gbx-A
Nedx
NK3
NK4-A
Repo
NK7-A
Lbx-A
Tlx-A
Pax4/6-2
Pax4/6-1
Drgx-B
Dmbx-A
Pitx-B
Dbx-A
7
Bari
prd-B
Emx1-A
Emx2-A
Pou6
Gbx-B-1
Zfhx-B
8
Ro-B
Meis-B
Hmbox-A
9
lab-B
pb-B
Hox3-B
Dfd-B
Scr-B
ftz-B
Antp-B
Ubx-B
abd-A-B
Abd-B-B
Phox-B
Onecut-B
NK5-B
Prrx
Noto-B
Lhx6/8
NK2.2-B
Otx-B
Hmbox-B
10
Pou3-A
11
Pou3-B
SatB
Prop
Onecut-A
Cux-A
Bsx1
Cers
Irx1-B
Irx3-B
Prox
Lhx2/9-2-B
En-A
12
Zeb
un-1
CG11294
NK2.1
Drgx-A
Meox
Abox
Pitx-A
Six1/2-B
Six3/6-B
Uncx-A
Shox
Cux-B
13
Bsx2
Meis-A
Noto-A-2
Noto-like-2
Noto-like-1
Noto-A-1
Vsx
Pax4/6-3
Six1/2-A
Six4/5-A
NK6-A
Lhx1/5-A
Msx2-A
Six3/6-A
Evx
Pou4-B
Rax-B
14
Otx-B-pseudo
Isl
NK2.2-A
Irx3-A
Irx2-A
Irx1-A
Rax-A
Hbn
Otp
Uncx-B

H)

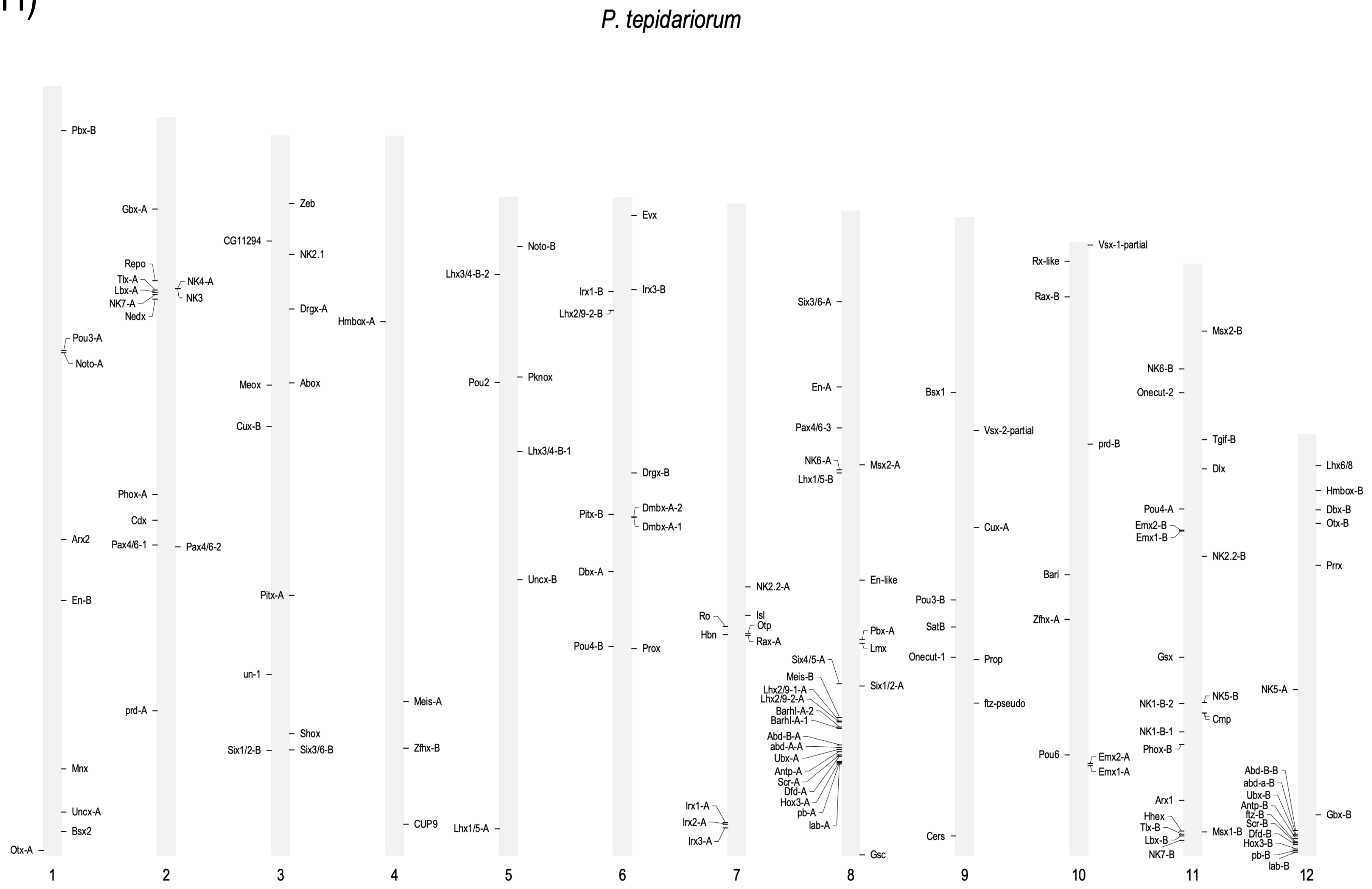
P. tepidariorum
1
Pbx-B
Pou3-A
Noto-A
Arx2
En-B
Mnx
Uncx-A
Bsx2
Otx-A
2
Gbx-A
Repo
Tlx-A
Lbx-A
NK7-A
Nedx
NK4-A
NK3
Phox-A
Cdx
Pax4/6-1
Pax4/6-2
prd-A
3
Zeb
CG11294
NK2.1
Drgx-A
Meox
Abox
Cux-B
Pitx-A
un-1
Shox
Six1/2-B
Six3/6-B
4
Hmbox-A
Meis-A
Zfhx-B
CUP9
5
Noto-B
Lhx3/4-B-2
Pou2
Pknox
Lhx3/4-B-1
Uncx-B
Lhx1/5-A
6
Evx
Irx1-B
Irx3-B
Lhx2/9-2-B
Drgx-B
Pitx-B
Dmbx-A-2
Dmbx-A-1
Dbx-A
Pou4-B
Prox
7
NK2.2-A
Ro
Isl
Otp
Hbn
Rax-A
Irx1-A
Irx2-A
Irx3-A
8
Six3/6-A
En-A
Pax4/6-3
NK6-A
Msx2-A
Lhx1/5-B
En-like
Pbx-A
Lmx
Six4/5-A
Meis-B
Six1/2-A
Lhx2/9-1-A
Lhx2/9-2-A
Barhl-A-2
Barhl-A-1
Abd-B-A
abd-A-A
Ubx-A
Antp-A
Scr-A
Dfd-A
Hox3-A
pb-A
lab-A
Gsc
9
Bsx1
Vsx-2-partial
Cux-A
Pou3-B
SatB
Onecut-1
Prop
ftz-pseudo
Cers
10
Vsx-1-partial
Rx-like
Rax-B
prd-B
Bari
Zfhx-A
Pou6
Emx2-A
Emx1-A
11
Msx2-B
NK6-B
Onecut-2
Tgif-B
Dlx
Pou4-A
Emx2-B
Emx1-B
NK2.2-B
Gsx
NK1-B-2
NK5-B
Cmp
NK1-B-1
Phox-B
Arx1
Hhex
Tlx-B
Lbx-B
NK7-B
Msx1-B
12
Lhx6/8
Hmbox-B
Dbx-B
Otx-B
Prrx
NK5-A
Abd-B-B
abd-a-B
Ubx-B
Antp-B
ftz-B
Scr-B
Dfd-B
Hox3-B
pb-B
lab-B
Gbx-B

l)

*H. graminicola*

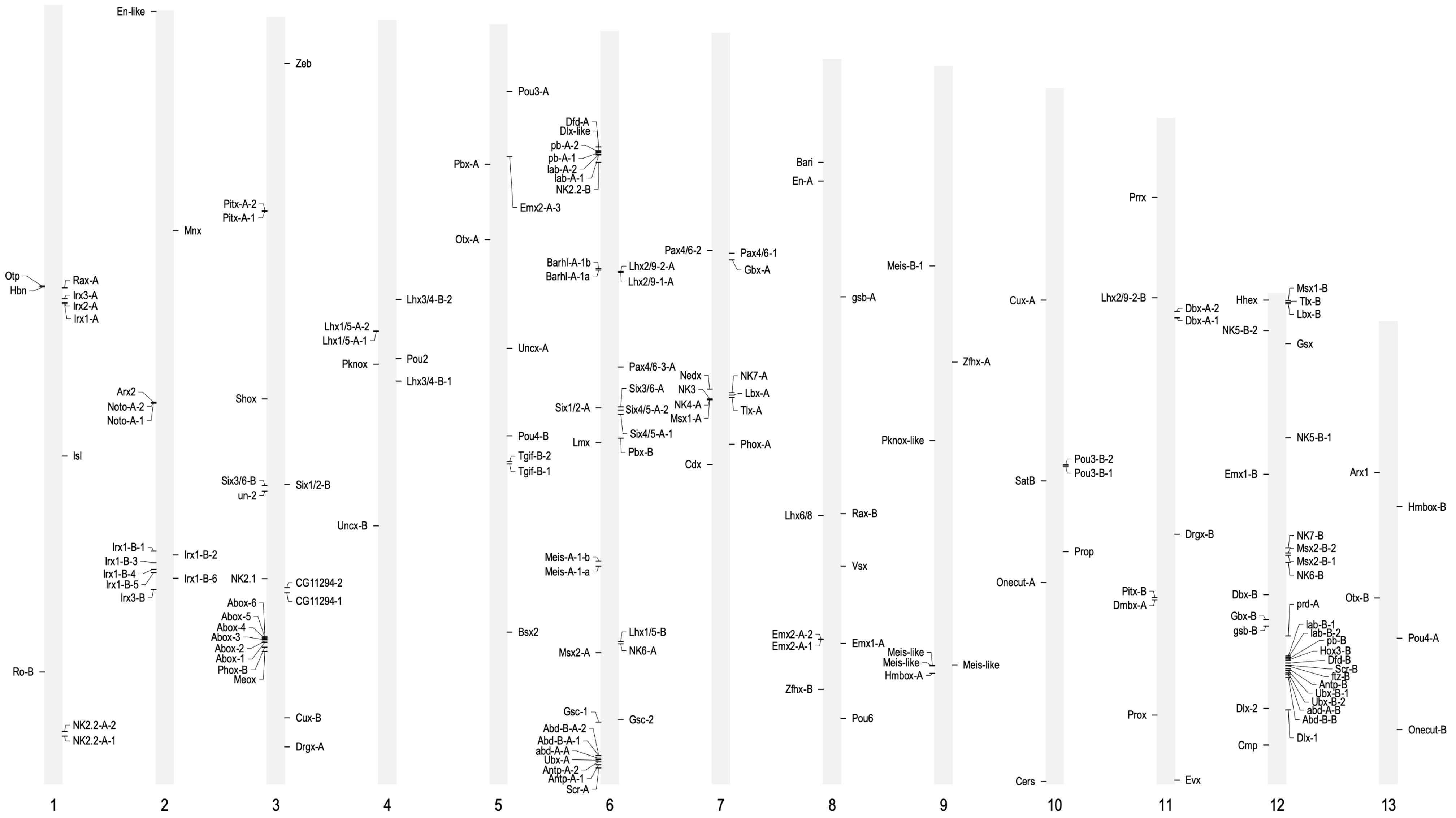

J)

*A. bruennichi*

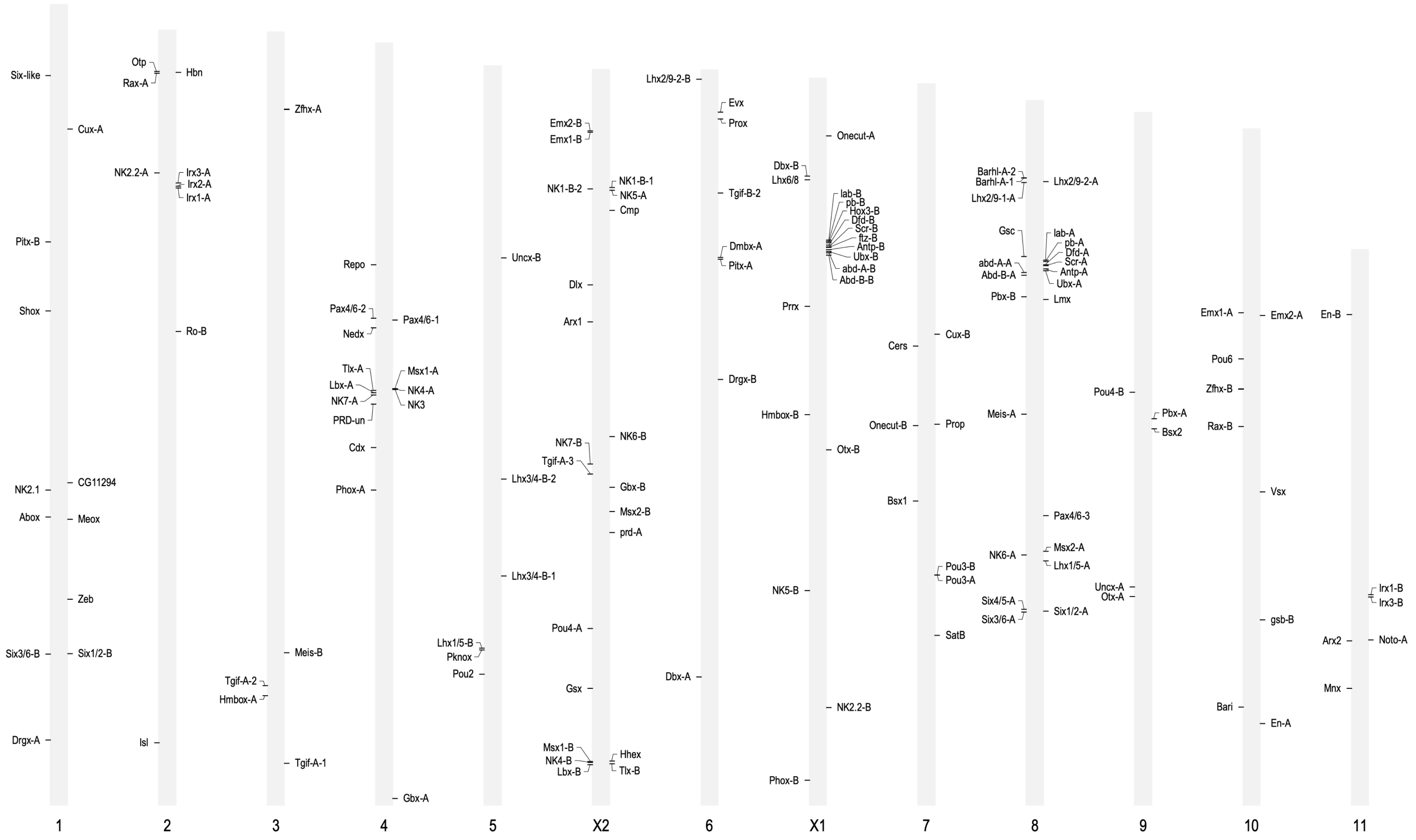

K)

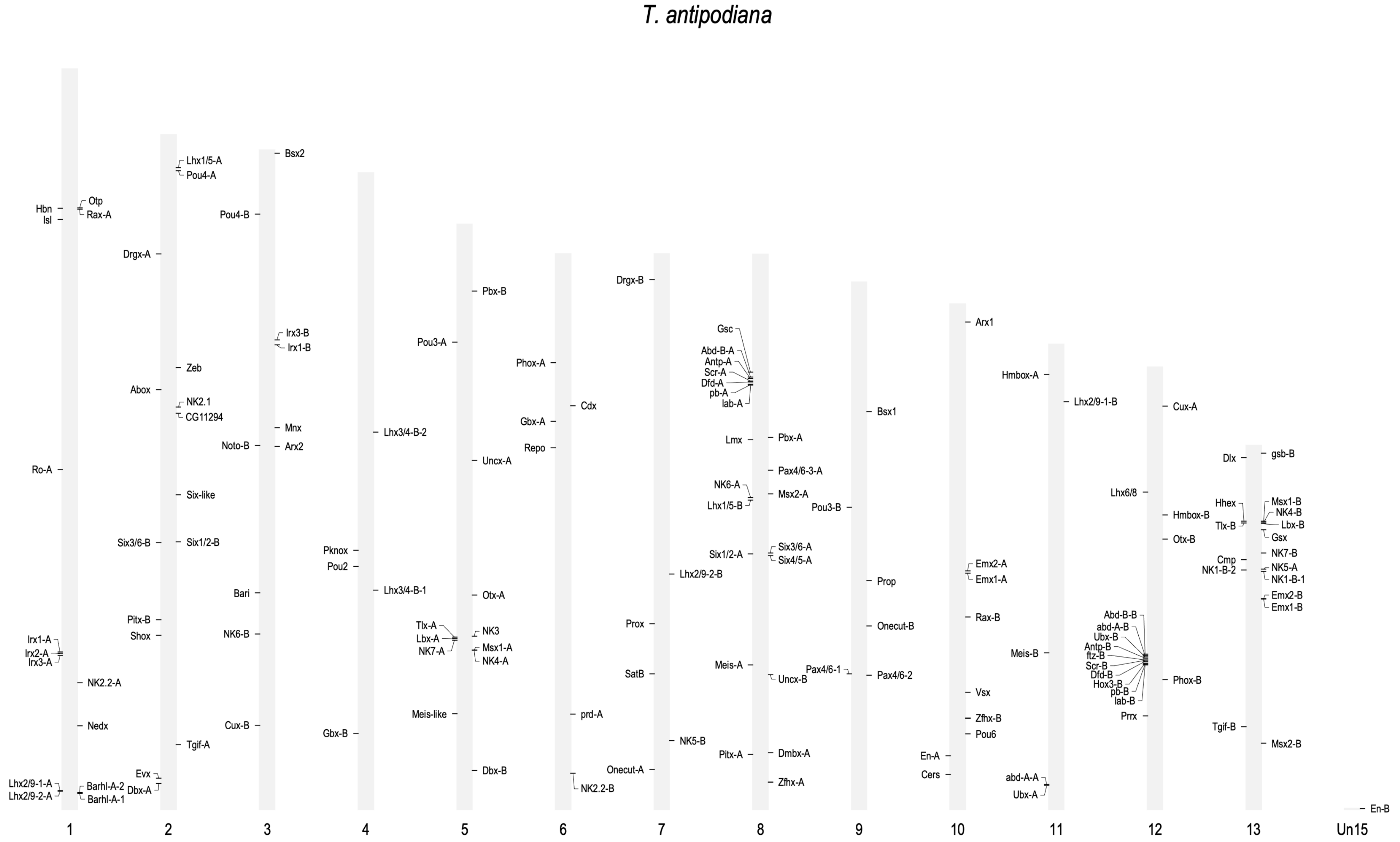
T. antipodiana
Hbn
Isl
Otp
Rax-A
Ro-A
Irx1-A
Irx2-A
Irx3-A
NK2.2-A
Nedx
Lhx2/9-1-A
Lhx2/9-2-A
Barhl-A-2
Barhl-A-1
Lhx1/5-A
Pou4-A
Drgx-A
Zeb
Abox
NK2.1
CG11294
Six-like
Six3/6-B
Six1/2-B
Pitx-B
Shox
Tgif-A
Evx
Dbx-A
Bsx2
Pou4-B
Irx3-B
Irx1-B
Mnx
Noto-B
Arx2
Bari
NK6-B
Cux-B
Lhx3/4-B-2
Pknox
Pou2
Lhx3/4-B-1
Gbx-B
Pbx-B
Pou3-A
Uncx-A
Otx-A
Tlx-A
Lbx-A
NK7-A
NK3
Msx1-A
NK4-A
Meis-like
Dbx-B
Phox-A
Cdx
Gbx-A
Repo
prd-A
NK2.2-B
Drgx-B
Lhx2/9-2-B
Prox
SatB
NK5-B
Onecut-A
Gsc
Abd-B-A
Antp-A
Scr-A
Dfd-A
pb-A
lab-A
Lmx
Pbx-A
Pax4/6-3-A
NK6-A
Msx2-A
Lhx1/5-B
Six1/2-A
Six3/6-A
Six4/5-A
Meis-A
Uncx-B
Pitx-A
Dmbx-A
Zfhx-A
Bsx1
Pou3-B
Prop
Onecut-B
Pax4/6-1
Pax4/6-2
Arx1
Emx2-A
Emx1-A
Rax-B
Vsx
Zfhx-B
Pou6
En-A
Cers
Hmbox-A
Lhx2/9-1-B
Meis-B
abd-A-A
Ubx-A
Cux-A
Lhx6/8
Hmbox-B
Otx-B
Abd-B-B
abd-A-B
Ubx-B
Antp-B
ftz-B
Scr-B
Dfd-B
Hox3-B
pb-B
lab-B
Phox-B
Prrx
Dlx
gsb-B
Hhex
Msx1-B
NK4-B
Tlx-B
Lbx-B
Gsx
NK7-B
Cmp
NK5-A
NK1-B-2
NK1-B-1
Emx2-B
Emx1-B
Tgif-B
Msx2-B
En-B
1
2
3
4
5
6
7
8
9
10
11
12
13
Un15

L)

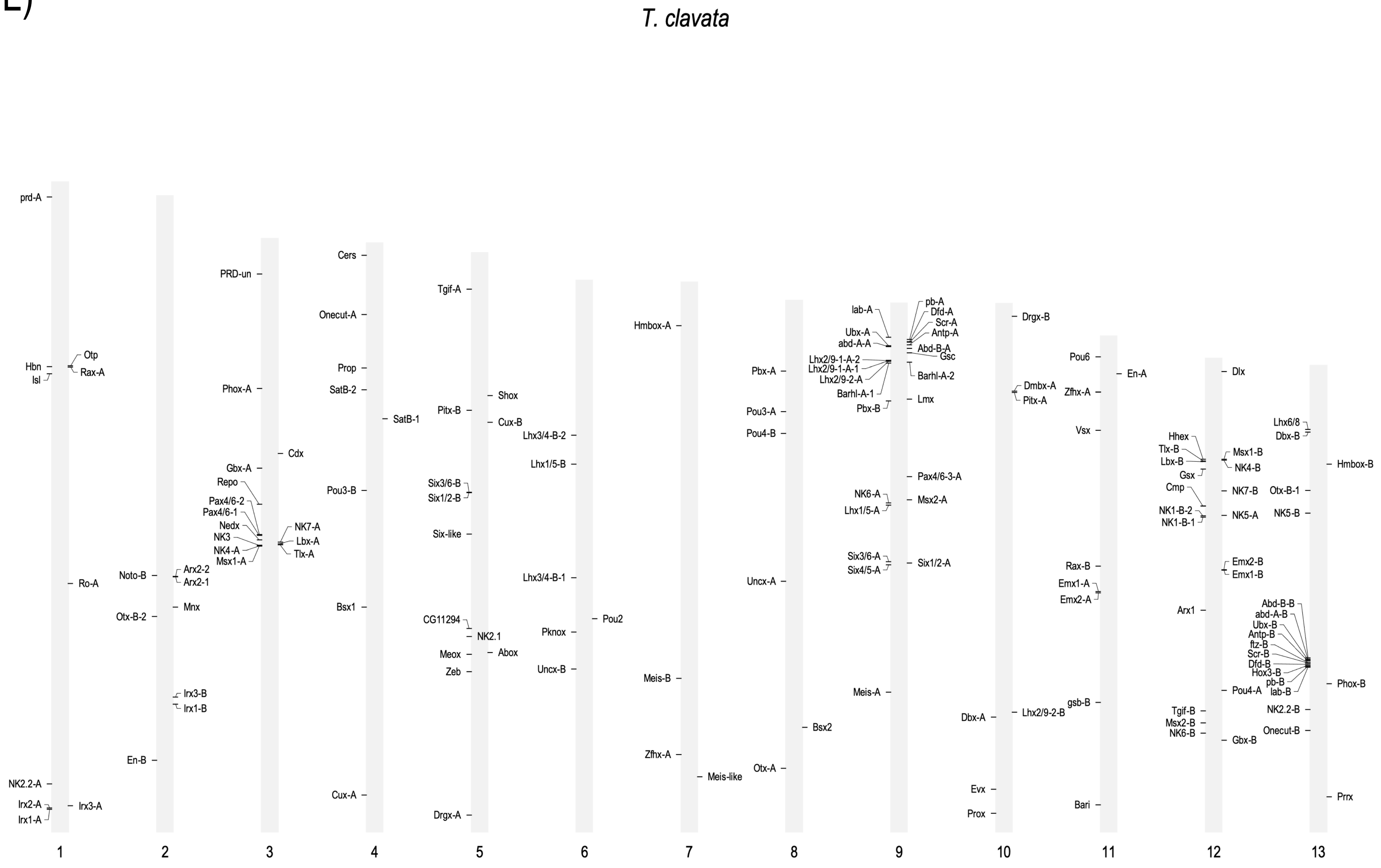
T. clavata
1
prd-A
Hbn
Isl
Otp
Rax-A
Ro-A
NK2.2-A
Irx2-A
Irx1-A
Irx3-A
2
Noto-B
Arx2-2
Arx2-1
Mnx
Otx-B-2
Irx3-B
Irx1-B
En-B
3
PRD-un
Phox-A
Cdx
Gbx-A
Repo
Pax4/6-2
Pax4/6-1
Nedx
NK3
NK4-A
Msx1-A
NK7-A
Lbx-A
Tlx-A
4
Cers
Onecut-A
Prop
SatB-2
SatB-1
Pou3-B
Bsx1
Cux-A
5
Tgif-A
Shox
Pitx-B
Cux-B
Six3/6-B
Six1/2-B
Six-like
CG11294
NK2.1
Meox
Abox
Zeb
Drgx-A
6
Lhx3/4-B-2
Lhx1/5-B
Lhx3/4-B-1
Pou2
Pknox
Uncx-B
7
Hmbox-A
Meis-B
Zfhx-A
Meis-like
8
Pbx-A
Pou3-A
Pou4-B
Uncx-A
Bsx2
Otx-A
9
lab-A
Ubx-A
abd-A-A
Lhx2/9-1-A-2
Lhx2/9-1-A-1
Lhx2/9-2-A
Barhl-A-1
Pbx-B
pb-A
Dfd-A
Scr-A
Antp-A
Abd-B-A
Gsc
Barhl-A-2
Lmx
Pax4/6-3-A
NK6-A
Msx2-A
Lhx1/5-A
Six3/6-A
Six4/5-A
Six1/2-A
Meis-A
10
Drgx-B
Dmbx-A
Pitx-A
Dbx-A
Lhx2/9-2-B
Evx
Prox
11
Pou6
En-A
Zfhx-A
Vsx
Rax-B
Emx1-A
Emx2-A
gsb-B
Bari
12
Dlx
Hhex
Tlx-B
Lbx-B
Msx1-B
NK4-B
Gsx
Cmp
NK7-B
NK1-B-2
NK1-B-1
NK5-A
Emx2-B
Emx1-B
Arx1
Pou4-A
Tgif-B
Msx2-B
NK6-B
Gbx-B
13
Lhx6/8
Dbx-B
Hmbox-B
Otx-B-1
NK5-B
Abd-B-B
abd-A-B
Ubx-B
Antp-B
ftz-B
Scr-B
Dfd-B
Hox3-B
pb-B
lab-B
Phox-B
NK2.2-B
Onecut-B
Prrx
