## Supplementary figures and images for "Evolution of the spider homeobox gene repertoire by tandem and whole genome duplication"

### FigureS1.pdf

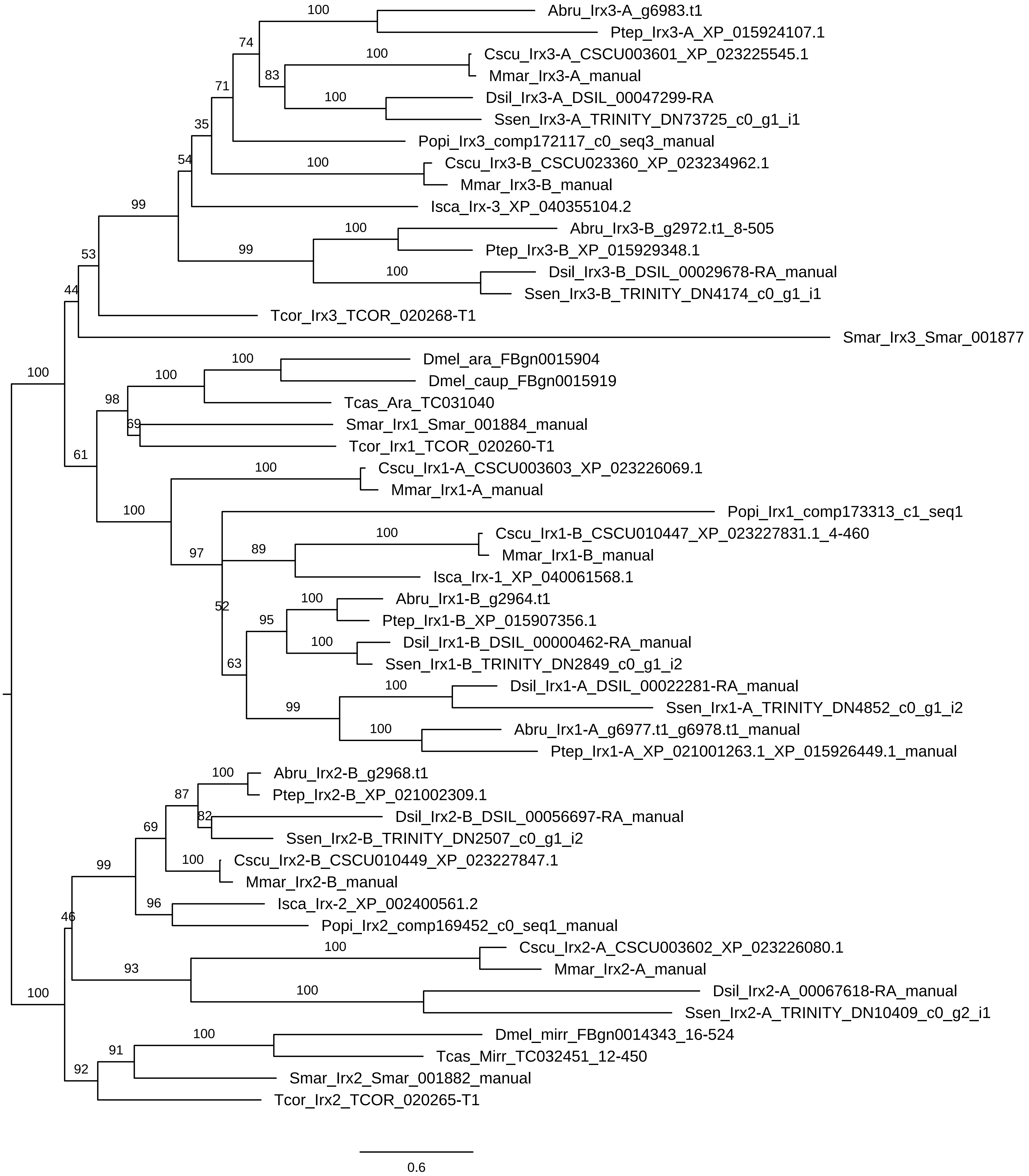

### FigureS2.pdf

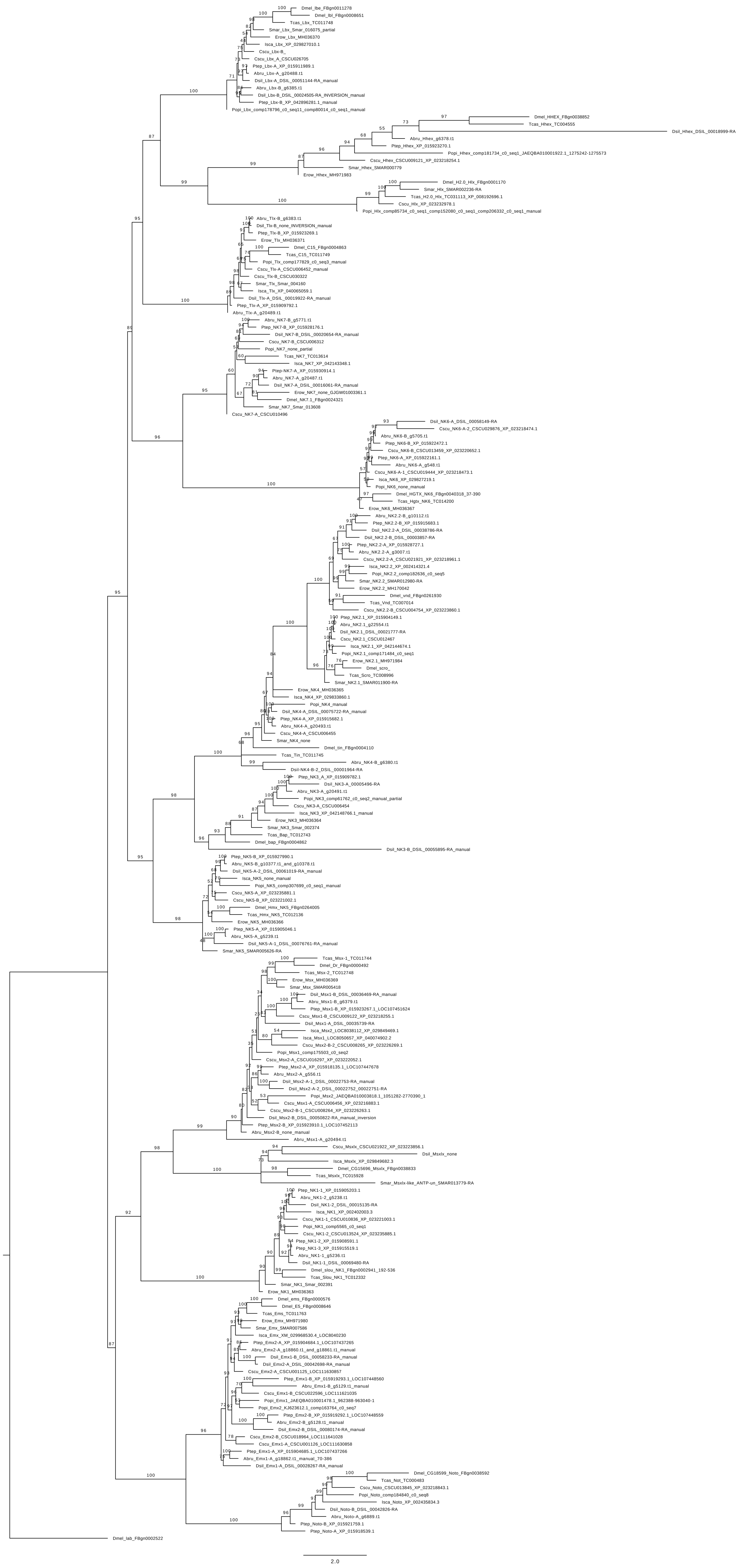

### FigureS4.pdf

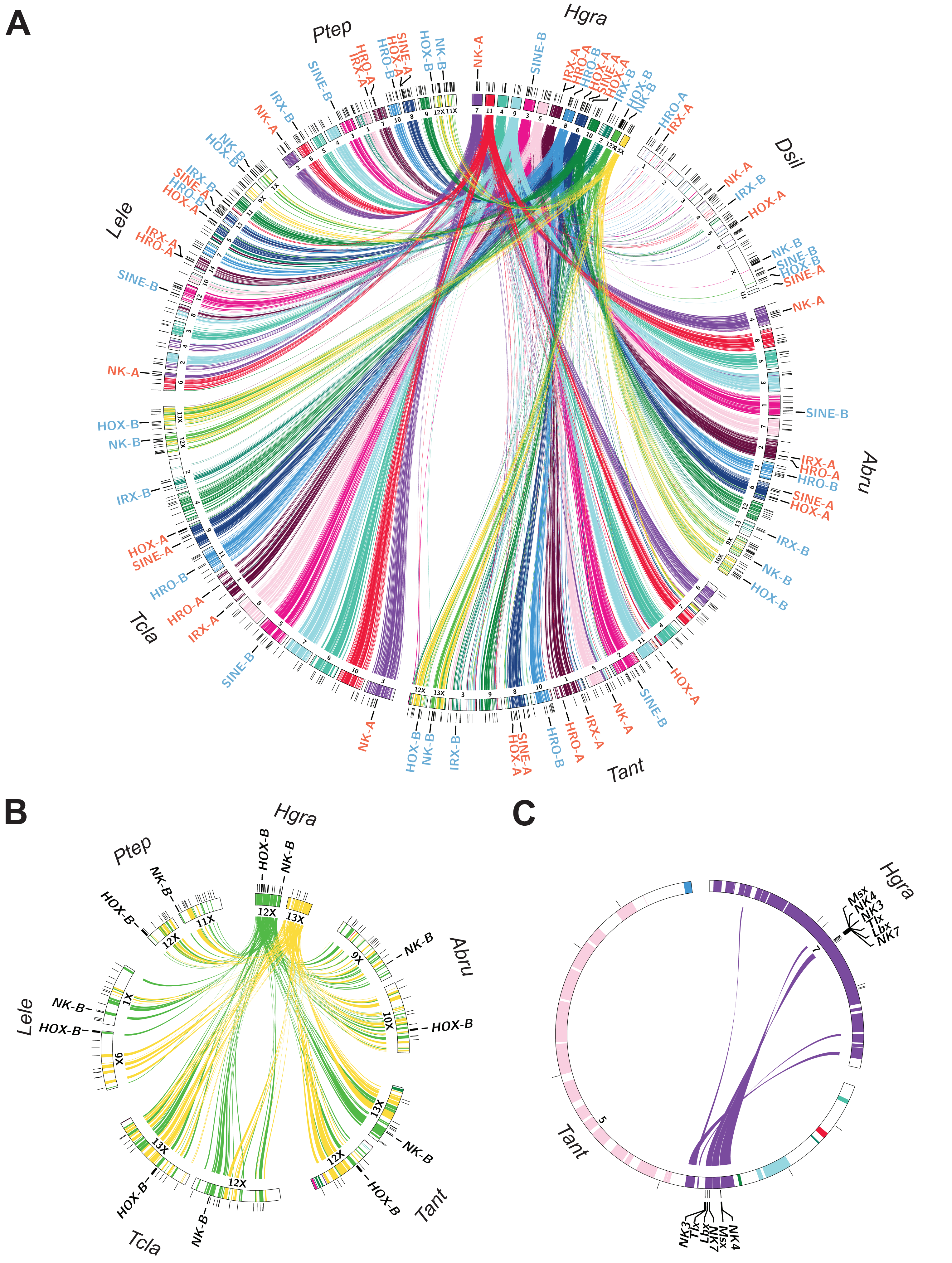

### FigureS5.pdf

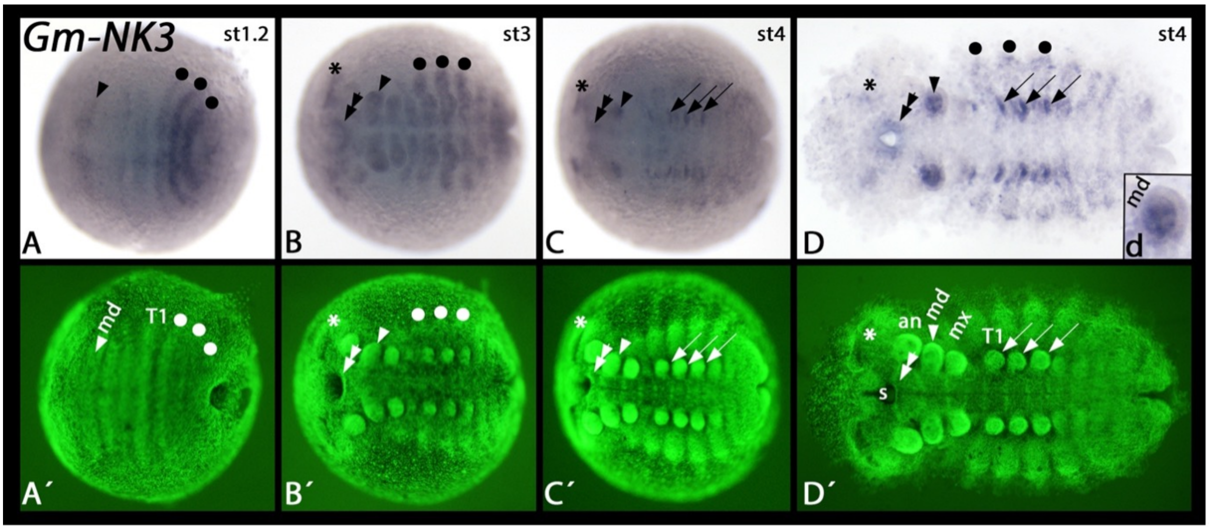
